# Why do we need high-fidelity synthetic eye movement data and how should they look like?

**DOI:** 10.64898/2025.12.11.692112

**Authors:** C. Stella Qian, Samantha Aziz, Kamrul Hasan, Oleg V. Komogortsev

## Abstract

Eye tracking has been a popular behavioral recording method across psychology, neuroscience, and computer science, but a need for large and diverse datasets has emerged. Synthetic eye movement data offer a promising complement, yet it remains unclear which aspects of real oculomotor behavior they must capture. This paper has three objectives: to clarify why synthetic eye movement data are needed, to outline what high-fidelity synthetic signals should look like, and to demonstrate how existing longitudinal datasets and subjective reports can guide their design and validation. We analyzed the motivation for synthetic eye movements and presented a framework of eye movement variance: ocassion-specific or state-specific variance, between-individual variance, pipeline induced variance and noise. Finally, we analyze subjective reports collected alongside the GazeBase dataset, demonstrating some ocassion-specific variance in data and setting requirements for state-free synthetic eye movement signals.

## 1 Introduction

Oculomotor data are laden with temporal information. Relative to other data which typically contain tens of hundreds or thousands of variables, eye movement data are rather simple: 2D coordinates of the gaze. All the other information is thus embedded in the temporal domain: simple eye movement components, gaze location and duration, eye movement as biometrics [1, 2], and eye movements as screening for certain disorders [3]. The intricate millisecond-level structure, including saccades, micro saccades, and fixation drifts, is one of the most popular and effective behavioral tools to draw insights into the neural processes of the brain on a wide range of topics such as perception [4, 5], attention [6, 7], memory [8], learning [9], and decision making [10]. Meanwhile, eye tracking devices sampling eye movements at hundreds of times per second are sufficient to provide rich data beyond the scrolling and clicks during web browsing. Unlike other typical behavioral interaction methodologies, eye tracking captures the underlying physiological and cognitive processes that drive the user’s behaviors, which can be useful for a large variety of applications. For example, eye movement data can support policymakers, companies, and consumers, by enabling better design and testing of policy and medical interventions, websites, store shelves, advertisements, and labels on food packaging [11]. Furthermore, eye tracking has been proven to serve as good biometrics and such identity-related features are stable enough across time. As a result, eye tracking is employed in academic research as well as product development, each leveraging it in various ways.

Oculomotor data, a rich source of information, has attracted interdisciplinary efforts in unraveling the mysteries. Such efforts often diverge in two major directions: a data-driven approach and a neuroscience theory driven approach. Here we hope to join forces on generating synthetic ocular motility data. For statisticians and computer scientists, the central challenge is to take advantage of the complexity of the system and generalize the functions into products or other functional packages. From the perspective of psychology, neuroscience, and biology, the key motivation lies in understanding human individual differences and cognitive processes underlying the eye movements while minimizing false positives and increasing replicability. Although these two approaches are dramatically different, there is one thing that can benefit both, namely, MORE DATA.

More oculomotor data with both quantity and quality can accelerate the progress of both fields. Data-driven models, particularly deep neural networks, are extremely data-hungry: large, high-resolution datasets improve generalization and mitigate overfitting. The model performance depends on data quality such as temporal fidelity, external noise, data loss in longitudinal datasets, and demographic coverage. For psychologists and neuroscientists, one challenge lies in obtaining sufficiently diverse samples outside the WEIRD (Western, Educated, Industrialized, Rich, and Democratic) data pool [12]. Thus, rich and diverse datasets not only support more robust statistical learning but also enable cross-population hypothesis testing, providing a shared foundation for both computational and theory-driven approaches. The obstacle to creating such datasets is that eye tracking data is expensive to collect, especially a large dataset.

Motivated by these challenges, we turn to the generation of synthetic oculomotor data as a complementary approach. Synthetic data offer a scalable and controllable way to explore hypotheses, benchmark models, and test algorithms under conditions that may be difficult or costly to produce experimentally. By simulating eye movement sequences with known ground truth, researchers can systematically vary task demands, noise levels, or participant characteristics and evaluate the robustness of their methods. While synthetic data cannot fully replace empirical recordings, they can accelerate model development and facilitate reproducibility by providing a shared, open resource for the community.

This paper encourages multidisciplinary and iterative endeavors toward generating synthetic oculor-motor data. Here, we describe the motivation, analyze the feasibility, and outline the methodological requirements. First, we aim to identify the requirements of future work and motivate that work on generating high-fidelity synthetic eye movement data by describing the complexities inherent in real oculomotor recordings, particularly those linked to user state changes during task performance, both temporary acute changes and the long-term changes. We then demonstrated how correlations emerge between objective measurements, features extracted from eye-tracking data, and subjective measures reported by participants; additionally, how these correlations fluctuate over time. To support transparency and future benchmarking, we publish the subjective data that we collected as a part of GazeBase, one of the largest and most widely used publicly available eye tracking dataset. In this paper, we do not aim to generate synthetic eye movement signals directly.

Finally, we curated a collaborative workflow for synthesizing eye movement signals (see figure 1): data collection, identifying the states of users or participants (e.g. fatigue), tasks, and resources of the data-driven approach and biological approach, conducting feature extraction and identifying variables of interest, discussing overlapping interests, generating synthetic signals, evaluating synthetic signals, and incorporating feedback into the next round of synthetic data generation. From the perspective of product development, relevant information is already available in the workflow. Specifically, the participants’ states, either characterized in subjective report form or calculated from behavioral data like reaction time, should demonstrate a robust correlation with features extracted from the eye movement data. This workflow is not limited to the synthetic generation of eye movements but extends to many other research domains that involve both physiological data and behavioral indices.

**Figure 1:**
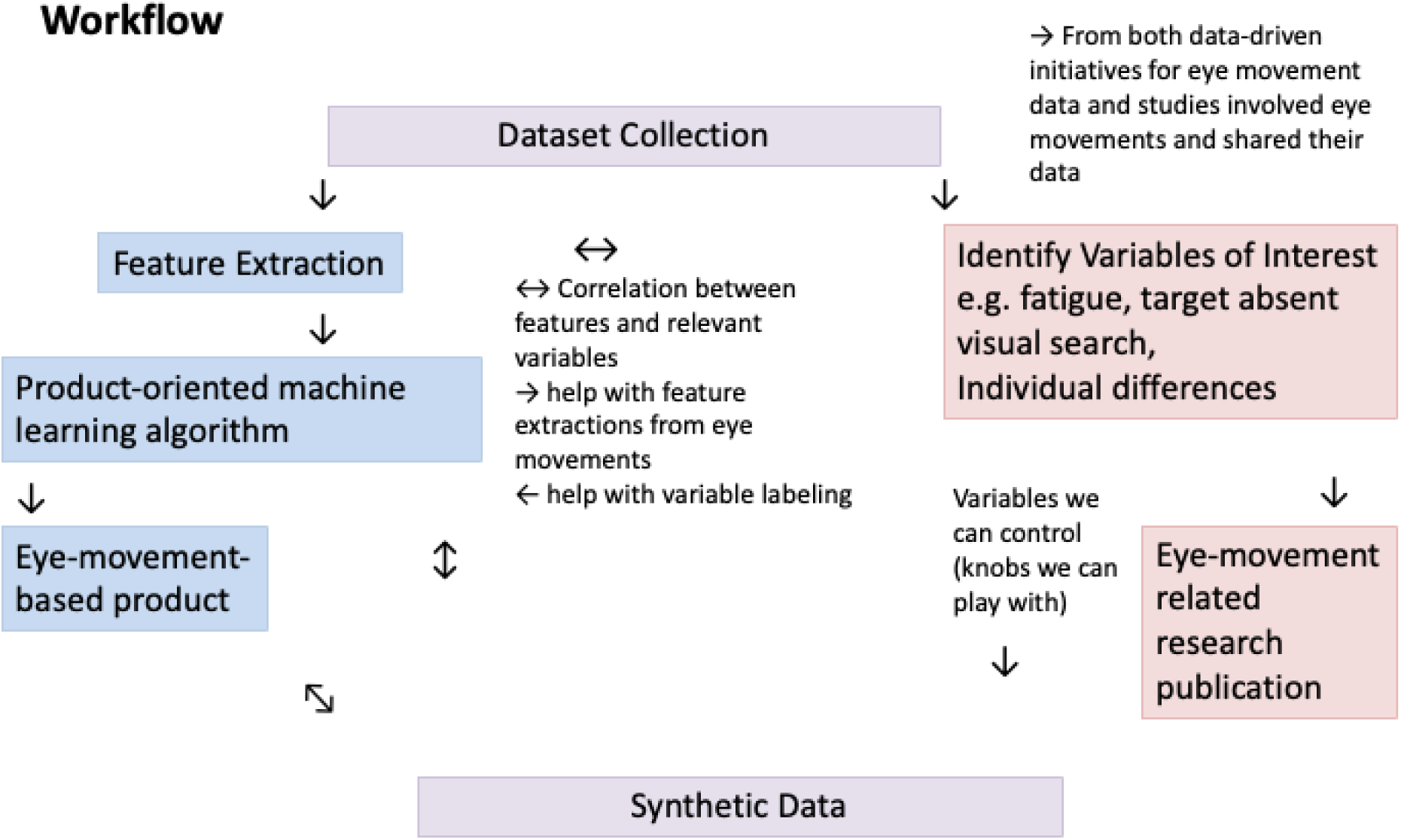
Workflow for synthesizing eye movement signals.

## 2 Motivation: The Need for Synthetic Eye Movement Data

### 2.1 Cost and Limitations of Real Data

Collecting high-quality eye movement data is an expensive process, often requiring significant financial, technical, and human labor investments. Eye-tracking equipment, particularly those offering high spatial and temporal resolution, is expensive, with costs ranging from several thousand to tens of thousands of dollars. In addition to hardware expenses, specialized software for data recording and analysis further adds to the financial burden. Moreover, proper experimental design demands controlled environments (e.g., light control), extended participant sessions, and trained personnel for calibration and supervision—each of which increases the overall cost. One of the largest high-fidelity longitudinal eye movement datasets is GazeBase, which consists of data collected using the EyeLink 1000 at 1000 Hz across 322 participants over 37 months [13]. Joint efforts were made on cross-linguistic eye movement recording during reading tasks: MECO recorded eye movement data during reading from 654 participants across 13 languages to improve cross-linguistic coverage in eye movement research [14]; MultiplEYE project is an ongoing international initiative to build a standardized, open multilingual reading eye-tracking dataset across dozens of languages with methodological consistency. Other notable high-resolution datasets include the ZuCo corpus, which combines eye-tracking and EEG data during natural reading tasks using an EyeLink 1000 Plus system [15], and DOVES dataset for natural image viewing using high-end desktop eye trackers [16]. While these datasets have fueled advancements in large scale eye movement research, these datasets may still be insufficient for computationally intensive applications such as deep neural networks, which typically require orders of magnitude more data to achieve generalization and robustness.

Beyond the high cost of collection, real eye movement datasets suffer from limitations that hinder scalability, diversity, and security. First, most datasets captured participants only once or twice, which makes studying long-term changes, such as biometric drift, fatigue patterns, or aging effects, very difficult [13]. Even datasets that achieved longitudinal data collection, such as GazeBase and GazeBaseVR, struggled with participant dropout (from 322 to 14 over 9 rounds in a 37-month period for GazeBase, 95.6% dropout rate; from 405 to 75 over 6 rounds in a 26-month period for GazeBaseVR, 81.5% dropout rate) and survey updating across different rounds of data collection [13, 17]. A second limitation arises from demographic and task homogeneity: most datasets recruit convenience samples of university students performing simple, artificial tasks such as a saccade task, reading, and passive viewing, with limited representation of ecologically valid activities like driving, gaming, web browsing, or mobile device use [16, 15]. This lack of diversity not only reduces generalizability but also biases models trained on such data. Privacy concerns further complicate the collection and sharing of real gaze data because eye movement traces can reveal sensitive personal biometric and cognitive information, leading to legal restrictions under frameworks like GDPR [18]. These problems are compounded by a lack of standardization: studies can substantially differ in eye tracking signal quality level at which data was recorded, calibration procedures, and event detection algorithms, which introduces noise and hampers reproducibility across datasets [19, 20]. Finally, gaze biometrics, although not globally recognized as a biometric property, but are still faced with the potential to spoofing. At this point, few public datasets are designed to test anti-spoofing methods, and raw gaze traces themselves are not intuitively interpretable to humans, making security issues less attended. Together, these limitations highlight the urgent need for high-fidelity synthetic eye movement data that can provide scalable, diverse, privacy-preserving, and standardized alternatives while capturing the rich temporal structure of natural oculomotor behavior.

### 2.2 Use Cases and Motivation

Resolving the scarcity of oculomotor data unlocks a broad array of applications. Imagine smart glasses with eye tracking becoming inexpensive enough to be affordable to the majority of human population, allowing one to wear a pair while reading. The ocular signals can serve as real-time indicators of cognitive states, labeling states such as active reading, momentary distraction due to boredom, comprehension struggles, or mind-wandering. These signals may also reflect fatigue, allowing the device to prompt beneficial rest reminders. Yet, the sensitivity of such data raises privacy concerns: readers may consent to share gaze data for training deep neural networks for reading comprehension tasks but they want their personal identity and medically related patterns private. Synthetic oculomotor data offers a solution: by generating realistic yet artificial eye-movement sequences, researchers can isolate and selectively reintegrate cognitive or affective factors into training without compromising privacy. Recent work such as SP-EyeGAN demonstrates how synthetic gaze signals can support reading comprehension models by enriching scarce datasets [21]. More generally, synthetic data approaches in healthcare and time-series analysis show how artificial datasets mitigate privacy risks and data scarcity problem, while retaining analytical utility [22, 23, 24]. Conceptually, synthetic gaze data function as a controlled tool for developing downstream training, where no real eye movement datasets can match.

Here, we invite researchers to join forces in studying eye movements by leveraging synthetic data generation. Unlike synthetic face images, the quality of synthetic eye movement data cannot be assessed by simple visual inspection. That is why it is important to summarize the key features of eye movement data in a single document. Similar to face stimuli, which are characterized by distinctive features like eyes and noses, eye movement patterns, eye movements are defined by their own fundamental components—blinks, saccades, pursuits, and so on. Each component occurs with its own logistics (e.g. we blink occasionally) and reflects the state or occasion. Meanwhile, the combination of these components can reveal the identity of the individual, a chronic mental state, temporal states (e.g. sleep deprived, high caffeine intake), and more. This paper hopes to delineate a structure for researchers who are interested in producing physiologically plausible synthetic eye movements.

### 2.3 Lessons from Other Domains

The use of artificially generated signals is best demonstrated with synthetic face images. These faces were typically generated by advanced AI models like GANs (Generative Adversarial Networks) [25], and they exhibited high realism, diversity, and controllability. These synthetic faces are crucial for testing and improving biometric systems, given the variety in texture, lighting, poses, expressions, and demographics[26, 27, 28, 29, 30, 31]. This is particularly helpful for understanding and correcting for biases: it was reported that the image quality of the synthetic faces was not consistent between racial and age groups [32]; StyleGAN models trained on the CelebA dataset produced faces with biases in skin tone, gender, and age due to under-representation [33]; fairFace datasets [34] and fairness-aware methods like conditional GANs [35] corrected these by balancing demographic attributes, improving biometric system fairness and informing face synthesis.

Generating synthetic faces share similarities with generating synthetic eye movement signals, as both involve generating complex, high-dimensional data to mimic human characteristics for biometric applications. Features in faces, such as the size of noses and the distance between eyes, are parallel to eye movement components like saccades and fixations. Both require algorithms to understand the variation of combinations of features. Both domains can also benefit from synthetic data to address privacy concerns and enhance dataset diversity, reducing biases in model training [35]. However, a key difference lies in the evaluation of the synthesized data: humans are inherently experts at interpreting faces but lack intuitive expertise in analyzing eye movement data. This demands interactive collaboration between researchers to re-evaluate synthetic eye movement signals every time new variables are involved.

Synthetic data have emerged as an essential resource across diverse research disciplines and practical applications, reflecting their broad utility and versatility. Examples include synthetic facial images generated by GAN for privacy-preserving facial recognition research [36, 37], artificially created medical imaging data to train diagnostic models without compromising patient confidentiality [38, 39], and synthetic text generated by advanced language models facilitate natural language processing tasks [40]. The primary motivations underlying synthetic data generation include preserving privacy, reducing costs and resource constraints associated with collecting real-world data, enhancing the diversity and scale of datasets to mitigate algorithmic bias, and enabling controlled experiments that precisely test specific hypotheses [22, 41]. Consequently, synthetic data have become ubiquitous, driving innovation and ensuring the responsible use of data across fields.

The success story from face synthesis offers insights into how to work on generating synthetic signals. Face synthesis researchers begin with large, diverse real datasets that include many identities, varied poses, lighting, expressions, etc., while ensuring careful annotations. Then, they built or trained generative models conditioned on the key variables of interest, such as pose, expression, illumination, occlusion, and identity. They also checked for distributional balance and latent disentanglement, and validate via downstream tasks (e.g. recognition) and via metrics on attributes. From these methods, we can learn how to define which components of eye movement (fixations, saccades, micro-saccades, pursuit drifts, temporal profiles), which state variables (e.g. cognitive load, fatigue, device noise), and which contexts (e.g. task type, participant variability) must be included, labeled carefully, varied broadly, and validated thoroughly.

The success story from face synthesis offers valuable insights into how to approach the generation of synthetic eye-movement signals. Face synthesis researchers begin with large, diverse real datasets containing many identities and systematically varied conditions such as pose, lighting, expression, and occlusion, while ensuring careful annotation of these variables. They then trained generative models, so that key variables like pose, expression, illumination, and identity can be explicitly controlled during synthesis [31, 36]. They worked to achieve distributional balance, ensuring that the dataset reflects a representative or deliberately controlled distribution of attributes (e.g., avoiding over-representation of frontal faces or certain demographic groups), which mitigates bias and improves generalization of face datasets[42]. They also aimed for latent disentanglement, structuring the generative model’s latent space so that changing one factor (e.g. expression) does not unintentionally alter others (such as identity), which enables controlled experiments and causal inference [43, 44]. Finally, they validate the realism and utility of the generated data through downstream tasks such as face recognition benchmarks and attribute classification, as well as human perceptual studies [45].

From these methods, we defined the key components of eye movement and their temporal profiles, as well as the temporal profiles of those components (Section 3). We proposed a few key variables, such as identity, fatigue, task difficulty, tasks, types of equipment, and more. These factors should be systematically labeled, varied in a balanced manner, and disentangled so that one can manipulate them independently of the others. Systematically labeling relevant attributes in the data allows more variables to be controlled in the training of the algorithm for the generation of the synthetic data. Such an approach will enable rigorous testing of models, causal investigations of oculomotor behavior, and fair benchmarking across diverse participants and contexts.

## 3 What Should Synthetic Eye Movement Data Look Like?

### 3.1 Oculomotor Components from a psychological and biological per-spective

Typical eye movement data contain a few distinct oculormotor events: blinks, fixations, saccades, and smooth pursuits. These elements are important as it is important for a face to contain eyes, noses, and mouths. And similarly, these oculormotor events have their own characteristic temporal signatures, interpersonal variability, and occasion-specific vulnerabilities. Spontaneous **blinks**, brief eyelid closures, occur at baseline rates of 12–20 blinks/min with mean durations of ∼250 ms; blink rate varies by ±30% across individuals and increases by ∼35% under mental fatigue (e.g., from ≈14 to ≈19 blinks/min after 2 h on task) [46, 47, 48]. **Fixations** are defined as periods of relative gaze stability when visual information is acquired. Typical fixations last on average 225 ms (range 50–1000 ms) and exhibit low within-task variability (±5 ms) but lengthen by ∼8% during increased lexical complexity (e.g., from 225 ms to 243 ms) [49, 50]. **Saccades** are the rapid ballistic shifts that reposition gaze and saccades usually have durations of 20–40 ms for small (*<* 5*^◦^*) amplitudes and latencies around 200 ms. Reaction times vary by up to 50 ms between individuals and slow by ∼20 ms under fatigue [51, 52, 53]. **Microsaccades**—small involuntary gaze shifts (*<* 1*^◦^* amplitude)—occur at rates of 1–3 per second with durations of 10–30 ms; although rates are relatively consistent, they can vary by ±0.5 per s across observers and increase during shifts in covert attention [54, 55]. **Smooth pursuit** is responsible for foveal tracking of moving targets and pursuits are initiated with latencies of ∼100 ms and achieve velocity gains near unity at target speeds ≤20*^◦^*/s, declining to ∼0.75 at 60*^◦^*; pursuit parameters are stable among healthy individuals but degrade by ∼10 % under divided attention [56, 57, 58, 59]. Finally, optokinetic nystagmus (**OKN**)—a reflex alternating slow-phase tracking and fast-phase resetting saccades in response to sustained visual motion—displays slow-phase gains around 0.8 with inter-subject variability <±0.05 but suffers ≥20 % gain reduction in cerebellar dysfunction [60, 61, 62, 63, 64].

Researchers in computational linguistics have made notable progress in simulating synthetic eye-movement data within reading contexts, notably through the simulation of fixation locations and durations tied to given texts [65]. However, many of these synthetic eye-movement models remain limited to reconstructing fixations and their durations, without recovering the full, temporally continuous high-fidelity oculormotor signals. For example, earlier approaches typically generate only aggregated fixation statistics or durations through relatively simple statistical models, which means they cannot support training or pre-training of advanced models that rely on raw, high-resolution scanpath data. While such gaze-only data are powerful for studying reading comprehension [66] and attention allocation [67], they are unlikely to be sufficient for applications that rely on individual-level identification or personalization, where subtle movement patterns across the full scanpath, such as microsaccades, are critical [65]. Furthermore, there is far more ocassion-specific information in the eye movements beyond fixations, which will be covered in the next section.

In contrast, generative models such as SP-EyeGAN can produce continuous sequences of gaze angles that resemble real human ocular micro- and macro-movements, enabling downstream applications like pre-training for reading-comprehension models. These methods go beyond reductionist gaze summaries and open the door to richer, more flexible synthetic data generation [21]. Diffgaze is a diffusion-based model that generates realistic continuous human gaze sequences on 360*^◦^* images, capturing temporal and spatial dependencies [68]. Same year, the same group proposed [69], a conditional diffusion-based framework that can generate subject-specific 5-second long realistic gaze sequences. Hasan et al. [70] recently analyzed and compared the GANs and the diffusion model for synthetic gaze sequence generation. Nonetheless, the earlier gaze-only models remain highly valuable for studying fixation-level behavior in reading. What we propose here is the development of more generic synthetic eye-movement generators that encompass not only fixations and their durations but also the whole spectrum of oculomotor dynamics, offering broader applicability across tasks while preserving the proven strengths of fixation-based simulations.

### 3.2 Oculormotor Components from a data-driven perspective

From a data-driven standpoint, oculomotor recordings can be conceptualized as the sum of multiple variance components arising from distinct sources. At the broadest level, the observed signal reflects a combination of person-specific variance—stable, identity-linked patterns in eye movement behavior [71, 72]—and state-dependent variance, which captures changes in that individual’s oculomotor behavior due to situational factors such as task demands, type of tasks, cognitive load, or environmental context, and residual noise pattern from a gaze estimation pipeline such as the recording devices and calibration protocols [73]. Beyond these explainable components lies a substantial portion of unexplained variance, often grouped under the umbrella term “noise.” This category may encompass a wide range of factors, from random fluctuations in neural control [74] to systematic biases introduced by the measurement process itself—such as differences between appearance-based gaze estimation methods and classical pupil–corneal reflection pipelines [75, 76, 77, 78]. Importantly, noise in this context is not necessarily devoid of structure; as shown in recent work [79], even high-frequency components traditionally treated as noise may retain meaningful biometric information. In this section, we will separately characterize inter-individual differences, task- or situation-specific variance, and noise-related components, providing both theoretical context and empirical findings for their identification.

#### 3.2.1 Occasion-specific and state-specific variance

Eye gaze is an important part of human social interaction. Our eyes move in patterned ways: human individuals all fixate, saccade, and pursue visual targets thousands of times per day—but beneath this uniformity lie subtle patterns. Even without eye tracking devices, observers visually examine others’ eye movement signals and judge their mental states, such as confusion, fatigue, or drug abuse. The majority of psychology research focusing on eye movements contributed to the understanding of occasion-specific or state-specific variance. In other words, what kind of consistent pattern do human observers produce when performing certain tasks or undergoing certain states. This section offers some examples from cognitive, emotional, aging, fatiguing, decision making, and psychopathological aspects, but these are far from the full spectrum of the occasion-specific and state-specific variance captured by eye movement signals.

##### Cognitive signatures

Cognitive signatures in eye movements have been extensively reviewed across multiple domains of psychology and neuroscience. Comprehensive reviews such as books [80, 81] that fixation duration, saccade dynamics, and pupil size act as sensitive markers of cognitive load, attention, and affective state. These syntheses highlight how eye-tracking provides non-invasive indices of momentary cognitive and emotional processes. Empirical studies support this perspective: Di Stasi et al. [82] found that saccadic peak velocity systematically decreases with rising mental workload, while Kaakinen and Hyönä [83] showed that task instructions modulate fixation durations and re-fixation probabilities during reading. Negi et al. [84] used an ecologically valid task to link fixation duration distributions to learning efficiency, demonstrating that longer and more variable fixations accompany deeper cognitive processing. Similarly, Katona et al. [85] observed that higher cognitive load elicited longer fixations, more numerous gaze shifts, and increased pupil dilation. Fadardi et al. [86] reported that elevated cognitive load slows saccade initiation and reduces corrective saccades, reflecting oculomotor inhibition under strain. Collectively, these findings confirm that eye-movement parameters encode observers’ cognitive states, making oculormotor dynamics a powerful window into transient fluctuations in attention and cognitive load.

##### Decision making

Eye-movement and pupillometric measures also carry rich signatures of decision formation in both perceptual and value-based choice. A robust finding is that gaze does not merely report preferences but also shapes them in a causal way. In binary value-based choices, the option that receives longer fixation time is more likely to be chosen, and this gaze bias is well captured by sequential sampling models in which evidence accumulation is multiplicatively boosted while an option is fixated [87, 88, 89]. Later work found evidence that looking or attention induced preferential choice [90]. Shimojo et al. [91] described a *gaze cascade* in which fixations progressively shift toward the ultimately chosen item during preference judgments. Subsequent work generalized these effects across domains, showing that individual differences in gaze bias parameters account for systematic variation in choice behavior [92]. These studies converge on the idea that moment-to-moment allocation of foveation is tightly coupled to the evolving decision variable.

Pupil diameter provides a complementary, arousal-linked channel through which decisional computations are expressed. Murphy et al. [93] showed that trial-to-trial fluctuations in baseline and task-evoked pupil size explain substantial variance in the speed and variability of perceptual decisions, consistent with models in which tonic arousal sets the gain of evidence accumulation. In a series of studies, Urai and colleagues demonstrated that pupil-linked arousal reflects decision uncertainty and shapes serial dependence in choice: larger post-decision pupil responses indexed higher model-derived uncertainty and predicted a greater tendency to alternate responses on the next trial [94]. Building on this, Urai et al. [95] used hierarchical drift–diffusion modeling to show that choice history biases subsequent decisions predominantly via changes in drift bias (how incoming evidence is interpreted) rather than starting point, indicating that history and arousal signals modulate the interpretation of current sensory input rather than only initial preferences. Converging evidence from de Gee et al. [96] shows that phasic pupil dilations are associated with a reduction in pre-existing decision biases across humans and mice, again via changes in accumulation bias, while Leong et al. [97] report that higher arousal can bias evidence accumulation toward motivationally desirable percepts. Collectively, these results suggest that pupil-linked arousal dynamically regulates how evidence is weighted, thereby tuning both the stability (history biases) and flexibility (bias suppression or motivational bias) of decision policies.

Finally, oculomotor kinematics, such as saccade vigor, provide an index of the latent decision variable and subjective value. Reppert et al. [98] demonstrated that saccades directed toward more highly valued options exhibit greater vigor (higher peak velocity for a given amplitude), implying that subjective economic value is partly expressed in the motor command. More recent work by Korbisch et al. [99] shows that saccade vigor systematically increases during deliberation and tracks the rise of the decision variable, with steeper increases when the value difference between options is larger. These findings dovetail with broader evidence that decision urgency and confidence modulate saccade metrics in speeded choice paradigms [100], supporting the view that the oculomotor system and decision circuitry are tightly co-regulated. Taken together, gaze allocation, pupil-linked arousal, and saccade dynamics jointly encode evolving decisional computations, providing a multi-dimensional, temporally precise window onto how humans evaluate options and commit to a choice.

##### Psychopathology

Abnormal eye movement signals have long been recognized as robust behavioral markers across several psychopathological conditions, with some of the strongest evidence emerging from schizophrenia research. Patients with schizophrenia show reliable impairments in smooth-pursuit eye movements, increased saccadic intrusions, and pronounced antisaccade deficits. Work by Thakkar and colleagues has clarified both the phenomenology and mechanisms of these abnormalities, demonstrating slower and fewer saccade error corrections [101], as well as systematic antisaccade impairments that track symptom severity [102]. Additionally, Friedman and co-workers found that smooth-pursuit performance in schizophrenia and affective disorders patients was significantly worse than in controls when quantified by RMS error and qualitative ratings [103, 104]. These deficits have been tied to disrupted cortical–subcortical circuits supporting prediction, inhibition, and sensorimotor integration, making them among the most replicable behavioral endophenotypes in schizophrenia. Similarly, atypical gaze patterns constitute a well-documented signature of autism spectrum disorder (ASD). Work by Klin, Jones and colleagues showed that autistic individuals preferentially fixate non-social objects over socially informative regions such as the eyes, and that reduced eye-looking predicts both clinical diagnosis and later symptom severity in infants at risk for ASD [105, 106]. Kevin A. Pelphrey et al. [107] demonstrated that autistic adolescents show distinct eye-movement patterns during biological motion perception, reflecting altered processing of socially relevant cues. Collectively, these studies establish social gaze atypicalities—reduced eye contact, altered fixation distribution, and atypical responses to social stimuli—as core oculomotor signatures of autism.

Attention-Deficit or Hyperactivity Disorder (ADHD) is associated with characteristic alterations in oculomotor control, reflecting deficits in sustained attention, inhibitory control, and sensorimotor regulation. One of the most robust findings comes from antisaccade and oculomotor inhibition paradigms: children and adults with ADHD exhibit increased antisaccade error rates and prolonged latencies, indicating impaired top-down suppression of reflexive gaze shifts[108, 109, 110, 111]. Smooth-pursuit deficits have also been repeatedly observed, with ADHD groups showing reduced pursuit gain, increased catch-up saccades, and less stable tracking [112], patterns interpreted as reflecting impairments in predictive control and visuomotor integration. Eye movement signatures also extend to behaviors in fixation: ADHD is associated with elevated microsaccade rates, greater gaze instability, and increased within-individual variability in fixation duration [108, 113], mirroring broader behavioral variability in the disorder. Together, these studies indicate that eye movement signatures, including antisaccade errors, pursuit gain reductions, fixation instability, and increased oculomotor variability, provide reliable, non-invasive indices of the attentional and inhibitory dysregulation central to ADHD.

##### Development

Beyond psychopathology, eye-movement behavior also changes systematically across the lifespan, reflecting the maturation and subsequent decline of the neural systems that govern oculomotor control. Many clinical studies—particularly in ADHD, autism, and schizophrenia—compare children and adults, underscoring the dramatic developmental improvements in fixation stability, saccadic precision, and inhibitory control that occur from childhood through young adulthood. Normative developmental work shows that children exhibit longer saccadic latencies, greater variability in saccade amplitude, and elevated antisaccade error rates compared with adults, consistent with the protracted maturation of fronto-striatal circuits supporting voluntary inhibition [114]. Smooth pursuit likewise becomes more accurate with age: pursuit gain increases steadily throughout adolescence as predictive mechanisms strengthen [114].

At the other end of the lifespan, healthy aging is accompanied by systematic changes in oculomotor signatures. Older adults consistently generate slower saccades with reduced peak velocity, increased latency, and longer duration saccades compared with younger individuals, reflecting age-related declines in sensorimotor execution and possibly brainstem burst-neuron function or cortical preparatory circuits [115]. Inhibitory control of eye movements also weakens with age: antisaccade and other oculomotor inhibition tasks show higher error rates, longer correction times, and greater variability in older adults than in younger cohorts [116]. Smooth-pursuit eye movements decline as well, with older participants exhibiting lower pursuit gain and more frequent corrective (catch-up) saccades, consistent with reduced motion-prediction capability and sensorimotor integration efficiency [117]. Moreover, fixation stability appears to degrade in late adulthood, with increased ocular drift and microsaccade frequency reported in aging samples, suggesting cumulative reductions in fine oculomotor calibration [118]. Together, these findings reveal that eye-movement metrics exhibit systematic, nonlinear change across the lifespan—improving through childhood into young adulthood and then declining in older age—making them a sensitive index of both developmental maturation and normative neural aging.

##### Fatigue

Fatigue is also found to be associated with eye movement signatures. Previous study showed a gradual slowing of saccade dynamics under fatigue, with some studies noting reduced saccade peak velocity, increased latency, or longer movement duration as individuals become more fatigued [119]. Fixation behavior may also be affected. Recent work using random-saccade paradigms has identified multiple fixation subtypes, and changes in the distribution of these fixation types have been observed during prolonged task performance [120]. Other signatures have been described as well: microsaccade rate often decreases while blink rate increases with fatigue, although the magnitude and consistency of these effects vary across tasks and participant samples [82]. Mental fatigue and eye tiredness were found to be different mechanisms according to an ERP study [121] and we need to dissociate them. Overall, while no single oculomotor metric provides a definitive indicator of fatigue, converging evidence suggests that saccades, fixations, microsaccades, and blinks each exhibit changes that may reflect accumulating mental or ocular fatigue over time.

##### Use Existing Datasets: Strengths and Gaps

Considering the extensive psychological and vision research on eye movements, existing datasets hold significant potential for reuse in developing and validating synthetic gaze data. Producing high-fidelity synthetic eye-movement signals requires access to diverse, high-quality recordings that span a wide range of tasks and participant backgrounds. While it is rarely feasible for a single dataset to meet all these criteria, combining data from multiple sources across disciplines offers a practical path toward comprehensive coverage.

Datasets collected by computer scientists or statisticians—such as GazeBase [13]—excel in scale and variety. They typically include large numbers of participants and a broad set of tasks, often with longitudinal recordings, enabling robust statistical modeling and cross-participant generalization. However, the tasks in such datasets are usually simple and brief (e.g., free-viewing or passive tasks with limited difficulty), which limits ecological validity and makes it challenging to study phenomena like extreme fatigue or motivation that emerge over extended, demanding sessions.

In contrast, datasets produced within psychology and vision science laboratories are often designed for hypothesis testing, featuring prolonged task durations, tightly controlled experimental manipulations, and rich metadata. These collections, though smaller in participant count (often 10–50 individuals), provide deep behavioral and physiological insights. Unfortunately, many of these datasets are difficult to locate or remain unpublished alongside the original research. They are often rediscovered only through review papers or secondary analyses, highlighting the fragmented accessibility of valuable eye-movement data.

When searching for existing data, researchers often rely on intuitive keywords such as “dataset,” yet this approach can overlook a vast array of publicly available eye-tracking resources embedded in supplementary materials or hosted on lab repositories. Despite the growing prevalence of confidence ratings and other behavioral measures in the literature, scientific progress has been slowed by the traditional lack of accessible, previously collected data. As a result, testing new ideas frequently requires researchers to spend months or years gathering data that, in many cases, already exist elsewhere. This inefficiency not only delays discovery but also discourages novel hypotheses that could be examined using the many underutilized datasets already produced by other scientists.

#### 3.2.2 Pipeline-induced Differences

The entire data-acquisition pipeline—from device selection through calibration, recording, and signal processing—can significantly affect the quality of eye-tracking data. As prior work has shown, both devices and methodological choices can systematically affect oculomotor data quality. For example, Holmqvist and colleagues [122] defined key metrics of data quality (accuracy, precision, and data loss) and outlined standardized procedures for their measurement across systems. Extending this work, other researchers have shown that methodological and environmental factors are important predictors of accuracy and data loss across commercial eye-trackers [123]. Recently, researchers have elaborated how hardware-and software-related choices (device type, head-free vs constrained setup, sampling rate), participant physiology (head-motion, eyes, calibration quality) and environmental conditions (lighting, distance) systematically predict accuracy and data-loss across commercial eye-trackers [73, 124, 125]. Other work demonstrated that latency (end-to-end delay) and precision differ markedly across even modern commercial systems: delays ranged around 15 to 52 ms, and latencies around 45 to 81 ms in HMD eye trackers [126]. Meanwhile, developmental and physiological factors show that precision and accuracy degrade in younger or less stable participants — indicating that the pipeline configuration (from optics, calibration, participant behaviour, recording to cleaning) can propagate error and bias into the recorded signal, making cross-device comparisons and reproducibility highly dependent on rigorous methodological control [127]. Taken together, these studies highlight that every stage of the eye-tracking pipeline—from optics and sampling to calibration and data cleaning—can propagate error and bias into the recorded signal, making cross-device comparisons and reproducibility highly dependent on rigorous methodological control. Researchers have already noticed this and collected data from two devices simultaneously to tackle the effect of device on eye tracking data [128]. Specifically, they recorded data with a wearable eye-tracking device (AdHawk MindLink eye tracker) along with a “gold standard” high-fidelity eye tracker (EyeLink 1000).

#### 3.2.3 Between-Individual Differences

Could eye movements give away information about the identity of individuals? We use faces or voices to judge people’s mental states but also their identities. Explicitly, we do not usually use another human’s eye movement patterns to recognize who they are. Neither do we use fingerprint or iris to identify identities but they both proved to be stable in individual characteristics. Eye movements also reflect biological and cognitive idiosyncrasies that vary systematically from one person to another. This raises a central question for both psychologists, biometric researchers, and computer scientists: are these differences large and consistent enough that we can reliably distinguish between individuals based solely on how their eyes move?

If we compare eye-movement signals to face images, which similarly encode both identity and states, we can gain insight into what makes biometric information interpretable and verifiable. Humans naturally rely on faces for identity recognition, which makes the distinctiveness of facial features intuitively obvious. In contrast, individuality in eye-movement patterns is not perceptually accessible; even trained researchers cannot visually identify a person from raw gaze traces. Like fingerprints, the discriminative information in eye-movement data is embedded in subtle, high-dimensional patterns that require computational or statistical analysis to detect. Consequently, establishing oculomotor signals as a biometric modality requires formal demonstrations and interpretive frameworks that explain how and why these latent features can encode identity.

Psychologists and biologists have long documented biologically influenced characteristics that persist within individuals over time in various oculomotor parameters. For instance, Ettinger et al. [129] assessed healthy adults on pro- and antisaccades and smooth pursuit across multiple sessions and reported high test–retest reliability (Intraclass Correlation Coefficients, ICCs = 0.77–0.85) for prosaccade latency (the time needed to initiate a rapid gaze shift toward a visual target; typically 190–220 ms), antisaccade error rate (frequency of mistakenly looking toward a target when instructed to look away; 8–15%), and pursuit gain (accuracy of eye tracking of moving objects; typically 0.70–1.00). Meyhöfer et al. similarly measured smooth pursuit at multiple target speeds (10, 20, and 30°/s) and reported ICCs greater than 0.70 for both pursuit gain and catch-up saccade frequency (0.2–1.5 saccades/s), confirming stable individual differences in pursuit accuracy [130]. Moreover, Klein and Fischer documented pronounced interpersonal variability (coefficients of variation around 20%) in the main-sequence slopes of peak saccade velocity—an established measure describing the relationship between saccadic amplitude and velocity (typically ranging from 200 to 600°/s)—highlighting consistent differences in eye-movement dynamics across individuals [131]. Smyrnis further observed stable individual variation in saccadic curvature (deviations of about 0.2 − 1.0*^◦^*, ICC ≈ 0.65), suggesting underlying anatomical or neural idiosyncrasies [132]. Genetic studies reinforce these findings, showing heritability estimates of roughly 40-50% for antisaccade errors and prosaccade latencies, indicating substantial genetic contributions to these measures [133]. Additionally, Vikesdal and Langaas demonstrated stable between-subject differences in pursuit latency (50–120 ms) and gain (0.60–0.90) across a wide range of target speeds (5 − 60*^◦^*/s), with reliability estimates between 0.60 and 0.80 [134]. Even ocular dominance asymmetries—where saccades directed toward the dominant eye are slightly larger (≈ 5*^◦^*) and faster—exhibit consistent yet subtle differences insufficiently robust for standalone individual classification [135]. Despite their reliability, individual oculomotor parameters generally offer moderate discriminability (e.g., Cohen’s d ≈ 0.5, an effect size also indicating substantial overlap between individuals’ scores). This moderate discriminability limits the effectiveness of single measures for uniquely distinguishing individuals [129, 131]. Therefore, combining multiple oculomotor features (e.g., latency, velocity, curvature, gain, and error metrics) into multivariate approaches is necessary to effectively differentiate between individuals or enhance clinical diagnostic accuracy [136, 137, 138, 139].

Researchers also found individual differences reflected in eye movements during reading. In particular, [140] reported stable between-subject variability across multiple eye-movement measures, including fixation durations and sensitivity to syntactic and semantic manipulations. Some readers consistently exhibited longer fixations and greater processing costs for complex sentence structures, whereas others showed faster, more efficient reading patterns with reduced disruption from the same linguistic challenges. Importantly, these differences were not random: the variability persisted across texts and conditions, suggesting trait-like properties similar to those documented for basic oculomotor parameters [130, 129, 131]. Moreover, complementary studies [49, 141] confirm that such variability in eye-movement metrics during reading is reliable over time and across tasks, reinforcing the idea that reading eye-movement data encodes robust individual difference information. These findings support efforts to integrate multiple oculomotor and cognitive measures to better capture the stable, person-specific signatures present in eye-tracking datasets [136, 142, 143].

From a computer science perspective, oculomotor signals present a valuable opportunity to model individual differences using high-dimensional, multi-featured representations. Whereas psychological and biological approaches often focus on a small number of interpretable traits, biometric methods in computer science can leverage a much broader set of features, mostly higher-order features, temporal dynamics, and nonlinear transformations—offering greater discriminative capacity. However, this advantage comes with a trade-off: many machine learning models in biometric research operate as “black boxes,” achieving high classification accuracy while providing limited transparency about which specific aspects of eye movements drive identification. Recent efforts have sought to bridge this gap by developing more interpretable feature representations. For example, Jäger et al. [144] characterized distributions of saccadic and microsaccadic features to capture meaningful variability, while Bolliger et al. [65] included features such as a word-order index to link sequential scan patterns. Together, such approaches suggest that interpretability and performance need not be mutually exclusive in modeling identity from eye movement signals.

From a computational modeling perspective, oculomotor plant models explicitly represent the physiological mechanisms underlying eye movements in anatomically and biomechanically plausible terms. Melnyk et al. [145] compared several classic and contemporary formulations—including Bahill’s foundational model [146], the updated version by Katrychuk et al. [147], and Enderle’s neurophysiological framework [148]—and demonstrated that Bahill’s model could capture substantial inter-individual variability using only 18 interpretable free parameters. Their analysis revealed critical subject-specific signatures in parameters such as the inertial mass of the eyeballs, the force–velocity characteristics of the agonist and antagonist muscles, and muscle elasticity, emphasizing that even relatively simple biomechanical models can effectively encode individual differences in oculomotor control.

Early demonstrations of eye-tracking-based biometrics by Kasprowski and Ober [2] showed that even simple features from reading and image-viewing tasks could identify individuals with a 1% false acceptance rate and 23% false rejection rate in a small group of 9 participants. Building on this, Komogortsev and colleagues systematically explored different families of eye-movement features. Holland and Komogortsev [149] examined a set of high-level average and aggregate eye-movement features, including fixation count, average fixation duration, average saccade amplitude and (peak) velocity, using a Gaussian distance function with weighted-mean fusion, they obtained an equal error rate (EER) of 22% in a subject pool of 32 participants. Komogortsev et al. [72] then introduced Oculomotor Plant Characteristics reviewed above, using mathematical models of the oculomotor plant to extract anatomical parameters of the human visual system from observable eye movements. In a study with 59 participants, this approach achieved a minimum half-total error rate of 19%. In related work, Komogortsev and Holland [150] focused on corrective oculomotor behavior, quantifying multiple types of saccadic dysmetria and express saccades and fusing information using likelihood-ratio tests, linear SVMs, and random forests, yielding an EER of 25% and a rank-1 identification rate of 47% for 32 participants. Then, Holland and Komogortsev [71] proposed low-level features based on the distributions of basic eye movements throughout a recording; comparing fixation and saccade distributions using the two-sample Cramér–von Mises test and fusing decisions with a 50-tree random forest, they reported a 17% EER and 83% rank-1 identification rate for 32 subjects.

Subsequent studies have demonstrated the scalability with deep neural networks. For example, Eye know you too used a denseNet-based conventional neural network and 5-second of eye movement recordings samples of 322 subjects, achieving state-of-the-art identification with an EER of just 3.66%, definitely acheiving a satisfying the FIDO Biometrics Requirements’ recommendation [1]. In parallel, the GazeBaseVR dataset enabled rigorous cross-session evaluation with consumer-grade VR headsets: using up to six sessions per participant across 407 individuals, researchers achieved binocular EERs as low as 1.67% and verified users with a false rejection rate of 22.73% at 10^-4 FAR [79], supporting the feasibility of scalable, session-robust biometric systems using immersive technology.

More ecologically valid, real-world settings have also been explored. In a wearable eye-tracking study of 39 individuals navigating familiar urban environments, [151] extracted five feature sets from mobile gaze data and trained random-forest models that achieved 78% identification accuracy and 89% verification accuracy (EERs of 6.3–9.1%), showing that person-specific gaze patterns persist even in complex, uncontrolled environments. Meanwhile, [152] evaluated 24 participants using nonlinear descriptors such as the largest Lyapunov exponent, achieving 100% classification accuracy in split-half validation, though between-session generalization remained a challenge—highlighting the gap between short-term and longitudinal robustness.

Addressing this issue directly, [153] evaluated continuous authentication models trained on GazeBaseVR over a 26-month interval. Short-term models (same-session or same-week) achieved up to 97% identification accuracy, but performance degraded sharply to as low as 1.78% over the multi-year span, unless adaptive retraining was employed—demonstrating the importance of temporal stability in practical biometric systems.

Together, these studies establish that eye-movement patterns, when modeled through rich, multi-variate computational techniques, contain substantial information that can be harnessed for reliable authentication. While challenges remain in interpretability, session generalization, and real-world deployment, it is now well demonstrated that eye movements encode sufficient individualized information to serve as an effective biometric modality [154].

#### 3.2.4 Unexplained variance: noise and beyond

A persistent challenge in oculomotor research lies in accounting for the unexplained variance within eye movement data, whose variance that does not neatly align with known signal components or behavioral phenomena. Multiple sources contribute to this unexplained variance: instrumental limitations (e.g., sampling rate, calibration drift, crosstalk) [155], physiological noise (such as microsaccades, eyelid interference, or physiological artifacts like heartbeat linked fluctuations [156, 157, 158]), and intrinsic neural variability within oculomotor control circuits [74]. Importantly, what we often label as “noise” may still convey identity-specific information. Raju et al.[79] examined this in Signal vs Noise in Eye-tracking Data: Biometric Implications and Identity Information Across Frequencies, demonstrating that while frequencies below ∼75 Hz largely constitute “signal”—carrying substantial biometric identity information—higher-frequency “noise” components also unexpectedly retain identity-specific patterns, even across both short-term (∼20 min) and long-term (∼1 year) intervals [159]. Further research will undoubtedly shed light on how much of this apparent “noise” can be reclassified as meaningful signal, thereby improving both interpretability and application in cognitive and biometric domains. Ultimately, the complex and multilayered nature of oculomotor data defies a simple binary classification, demanding further experimental work, dataset building, and analytical methods that recognize its potential as both a confound and a source of valuable information.

## 4 Analyses of Subjective Reports from GazeBase Dataset

Associating subjective data with objective features is critical for studying the meaning of eye movement data, as well as for product development. To facilitate these goals, we released further data from GazeBase (see Appendix for more information about GazeBase and all the data and code for this project is published in the Appendix). Subjective data are valuable sources of evaluative properties of a dataset that are expensive to collect, so that it is critical to measure them as accurately as possible. Here, we examined all the subjective report data from GazeBase dataset, evaluated the quality and characteristics of the subjective reports, associated these subjective reports with objective measurements, in this case, the eye movement data. We hope to offer insights into how synthetic eye movement data should look like and behave like, encourage both psychologists and computer scientists to explore this oculormotor dataset, and facilitate product development using this rich longitudinal dataset with subjective reports.

Participants performed subjective reports at various points during the experimental sessions. Their eye movements were recorded as participants performed a battery of tasks (the description of these tasks can be seen in this paper [13] as well as the appendix method section) with high-fidelity eye tracking. The battery of tasks were tested twice for each visit in the lab, which we call sessions. Subjective reports were obtained before sessions, between two sessions, and after sessions: participants reported general comfort, shoulder fatigue, neck fatigue, and eyes fatigue for every report, and physical effort and mental effort to complete tasks between- or after-session. The survey also collected further information such as caffeine intake, sleep, alcohol, and headache. These surveys were made publicly available along this manuscript. To further understand the effect from different tasks, subjective reports after each task were introduced in the data collection after the first round [13]. After each experimental task, the participants were asked about how difficult the task was, their mental tiredness, and their eye tiredness on a Likert scale 1-7 (1 being not tired or difficult at all and 7 being very tired or very difficult). We acknowledge the limitation of these measures, so we would like to evaluate how informative these subjective reports are and how consistent they are.

### 4.1 Reliability of subjective reports after each task

We performed descriptive statistical analyses and conducted correlation analysis between two sessions as an examination of the reliability of these measures, following by an exploratory factor analysis to see which measurements and which tasks tend to group together. These comprehensive analyses systematically evaluate the underlying structure of subjective reports obtained in between oculomotor experimental tasks, providing a robust methodological foundation for understanding these measurements and different tasks. Further, we examined one task that demonstrated the largest amount of variation in subjective reports, speculating on the meaning of such results.

The distribution of participants subjective report is shown in 2. Each plot is subjective ratings as a function of different task. The sessions and the reported items are presented in separate subplots. The ratings have a floor effect, indicating that the majority of the participants did not report to be fatigued in all three items in most of the tasks. A correlation analysis was conducted to assess the test-retest reliability of subjective measures between two experimental sessions (see Figures 3, 5, and 4). We assume that participants’ subjective ratings of the same task stay consistent on the same day within a short period of time, namely between two experimental sessions. We chose Pearson correlation to examine the subjective rating consistency between two sessions because. Pearson correlation coefficients were calculated for each subjective measure (subjective task difficulty, mental tiredness, and eye tiredness) across all experimental tasks to evaluate consistency over time. Test-retest reliability was considered adequate when correlation coefficients exceeded 0.70, following established psychometric standards.

**Figure 2:**
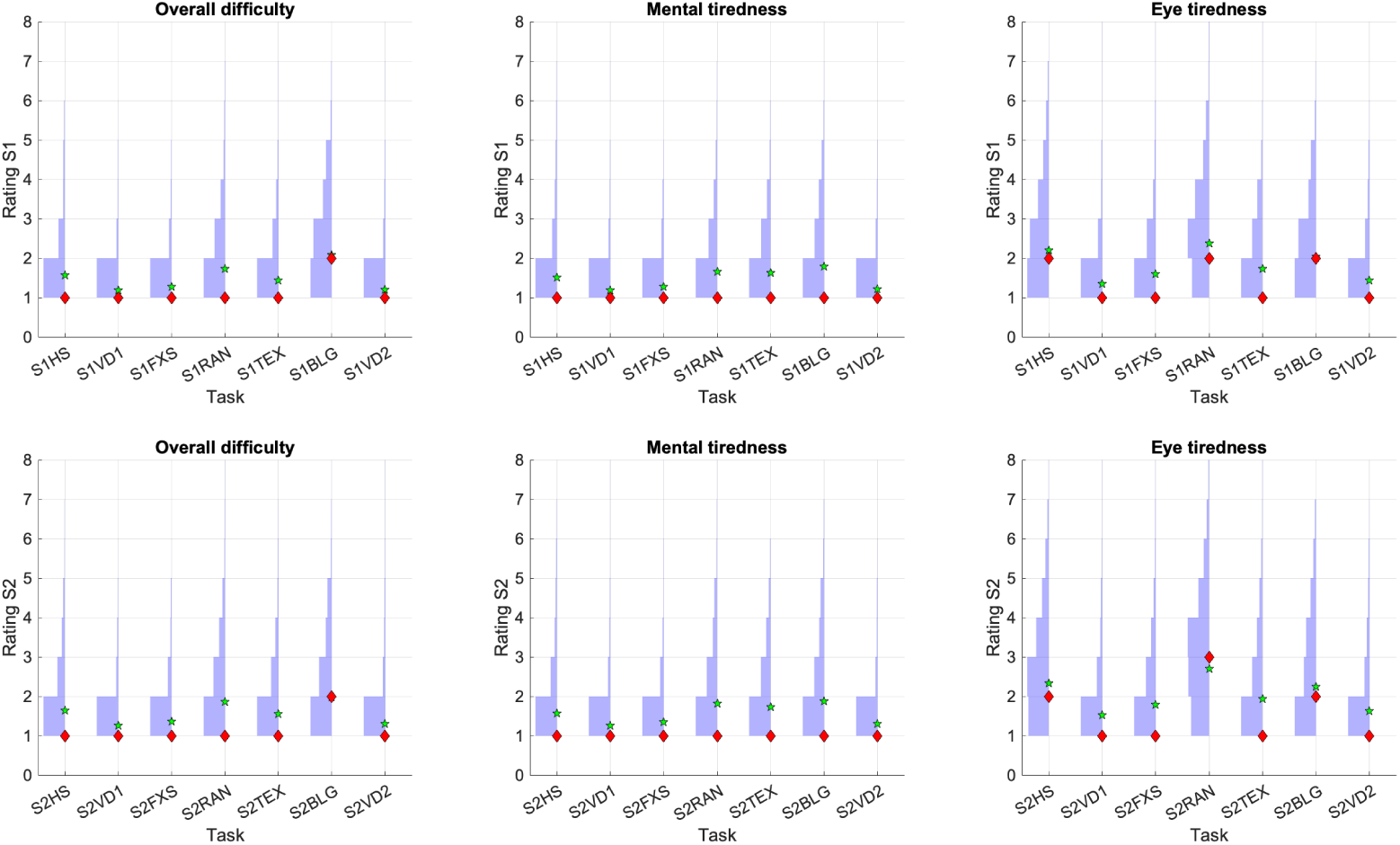
Distribution of subjective reports after each task. The two rows correspond to the two experimental sessions during each visit. Each column correspond to a specific item in the survey. Each subplot is transposed histograms as a function of tasks. The green star represents the mean of the subjective rating and the red diamond represents the median.

**Figure 3:**
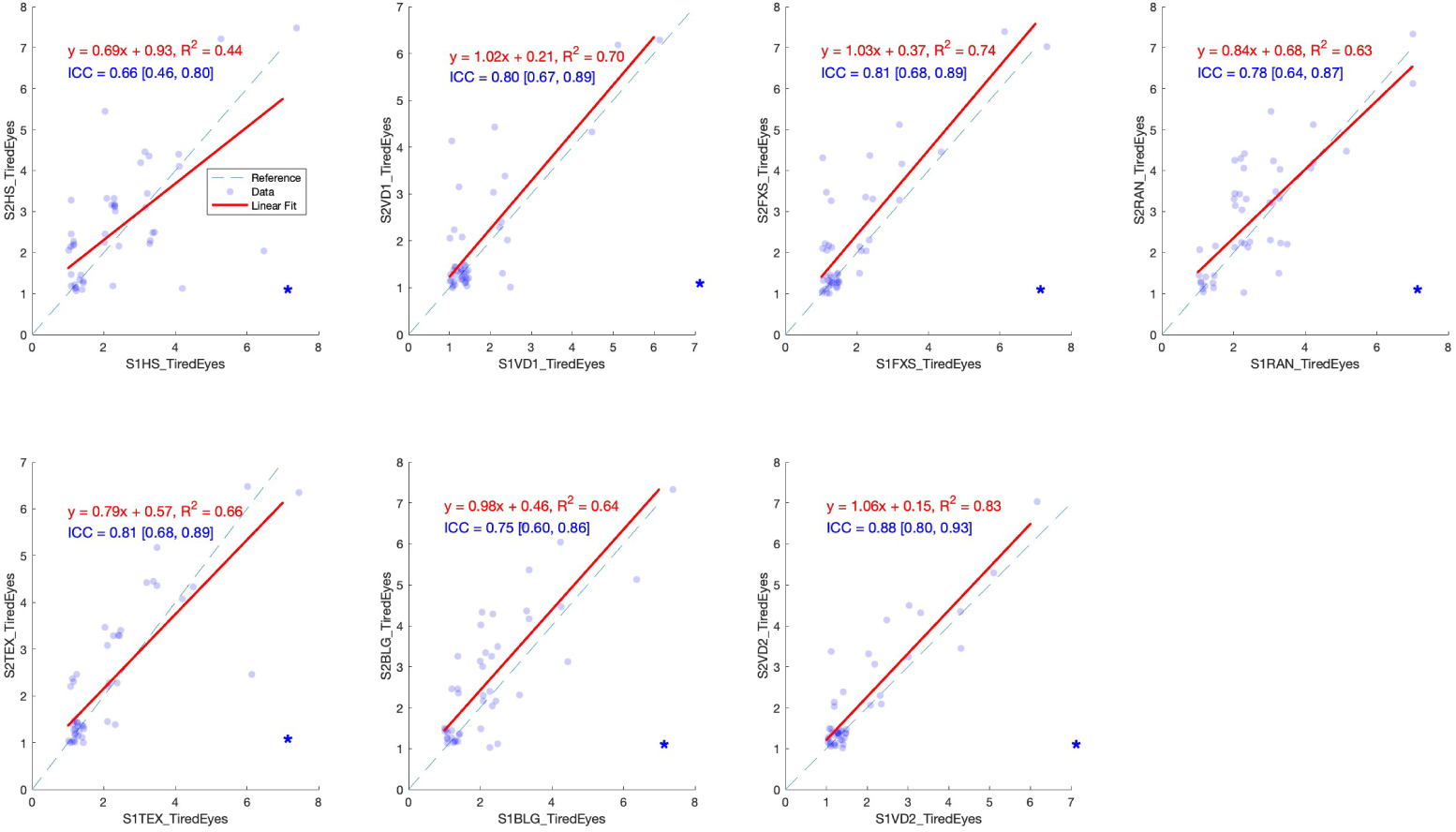
Correlation of ratings of eye tiredness between two sessions of participation. Each panel represents the correlation of a different task. The linear fit and the R 2 are denoted in the figure. For the best display of the data, the scatter plots representing each individual participants’ data were jittered around the integer, but only the raw data were used for analyses. The blue asterisk denotes statistical significance of correlation.

**Figure 4:**
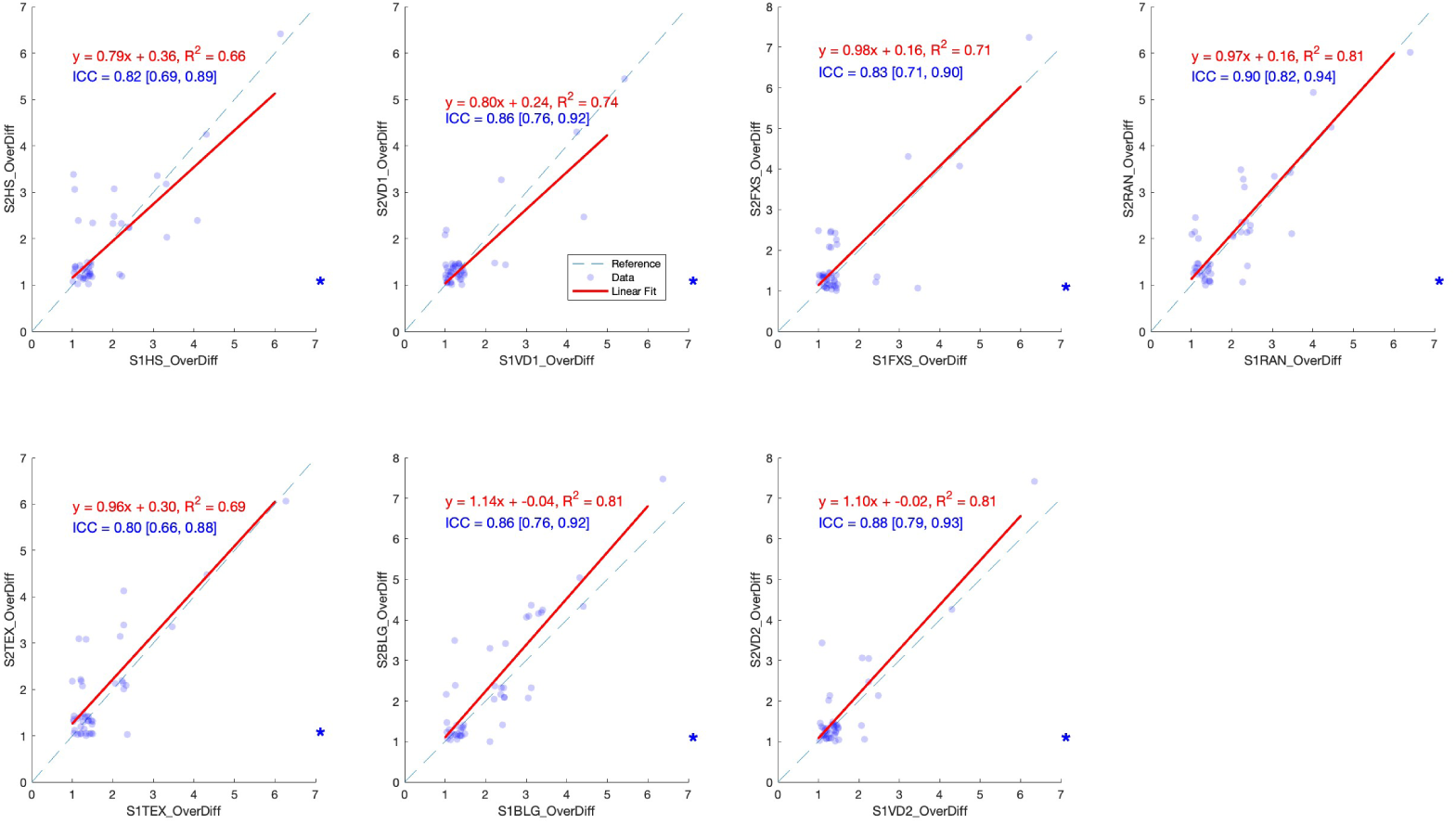
Correlation of ratings of overall difficulty between two sessions of participation

**Figure 5:**
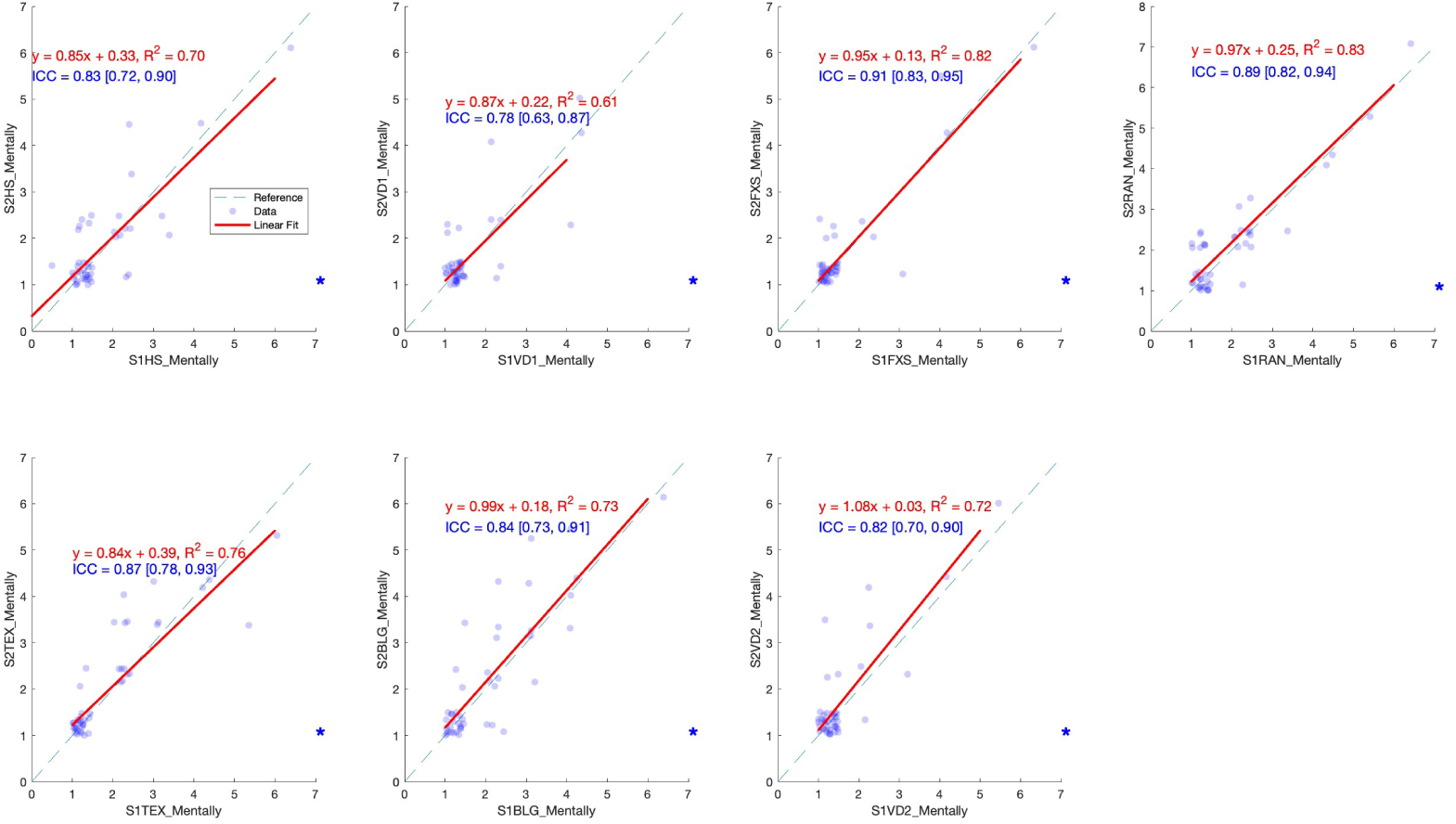
Correlation of ratings of mental tiredness between two sessions of participation

Test-retest reliability is a traditional methodology to examine whether a measurement is producing consistent results [160]. This dataset offered us an opportunity to do so by examining the correlation and intraclass correlation (ICC) between two experimental sessions every participant performed within 30 minutes. We conducted a linear regression between the ratings after the same task in two sessions, and participants did three ratings on mental tiredness, eye tiredness, and subjective difficulty. That resulted in 21 pairs (7 tasks and 3 ratings) of correlation analysis (see figure blahblah for raw data, linear fitting, and the fitting function). All of them produced significant correlation and ICC higher than 0.6 except for one pair (eye tiredness for VD1 task 0.44). Additionally, intraclass correlation coefficients (ICC) were calculated to provide a more robust measure of agreement between sessions, accounting for both correlation and systematic differences in ratings. ICC values were interpreted according to standard guidelines: values <0.50 indicating poor reliability, 0.50-0.75 indicating moderate reliability, 0.75-0.90 indicating good reliability, and >0.90 indicating excellent reliability[161]. This analysis allowed us to determine whether subjective reports maintained consistent patterns across experimental sessions, thus validating their utility as reliable indicators of participant experience.

### 4.2 Factor analysis of subjective reports after each task

An exploratory factor analysis (EFA) was performed to understand the underlying the latent structure of these batteries of tasks and subjective measures. This EFA used principal axis factoring with oblimin rotation was conducted on 42 items (7 tasks * 2 sessions * 3 questions). The KaiserMeyerOlkin (KMO) measure of sampling adequacy was .97, and Bartlett’s test of sphericity was significant (*χ*^2^(861) = 32702, p < .001), indicating that the data were suitable for factor analysis.

Five factors were extracted based on the scree plot (see Figure 6) and eigenvalues greater than 1, explaining 81.2% of the total variance. However, based on the interpretation of the meaning of the factors, we arrived at 3 factors, explaining 73.6% of the variance. The factor loadings can be found in Table 1. The interpretation of the factors were described below and the rotated factor solution is presented in Table 1. Items were considered to load onto a factor if they had a primary loading ≥.40 and did not cross-load above .30 on other factors.

- **Factor 1 (Viewing task factor):** Included items related to over-difficulty and mental tiredness on some passive viewing tasks (VD1, VD2, and FXS), such as S1VD1_OverDiff (.71), S1VD1_Mentally (.71), and S1FXS_OverDiff (.71).
- **Factor 2 (Saccade related factor):** Comprised mainly items measuring overall difficulty and mental tiredness from the saccade heavy tasks such as RAN, HS, and BLG. For example, S1RAN_OverDiff (.77) and S1RAN_Mentally (.76). The reading task has cross loading of both Factor 1 and Factor 2.
- **Factor 3 (Eye tiredness factor):** Characterized by tired eyes ratings across all different tasks (e.g., S1FXS_TiredEyes = .60; S1VD1_TiredEyes = .60). Multiple items have cross-loadings on other factors.

**Figure 6:**
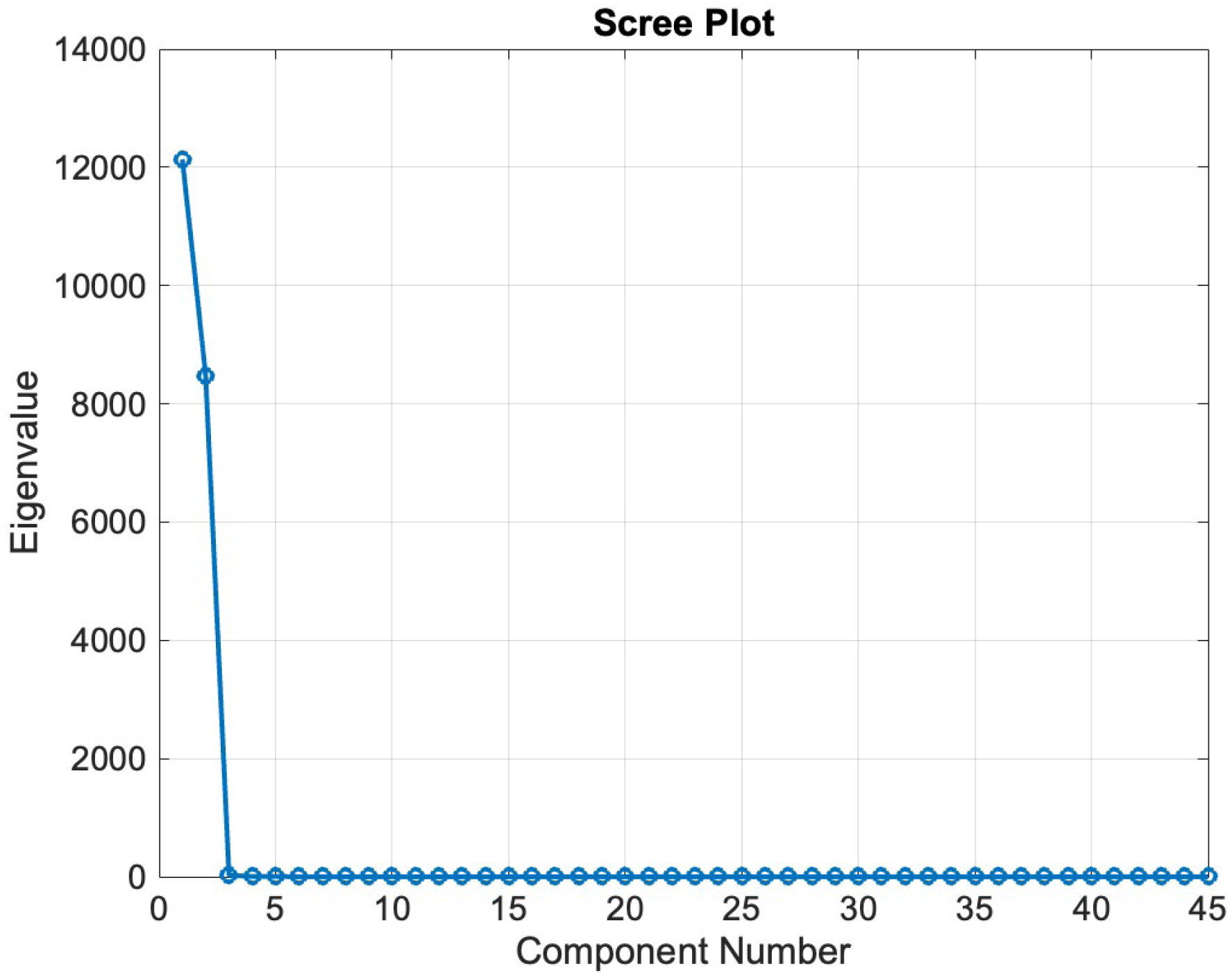
Scree plot of exploratory factor analysis

**Table 1:** Factor loadings of EFA from subjective reports data.

|  | Factor 1 | Factor 2 | Factor 3 |
| --- | --- | --- | --- |
| S1HS_OverDiff | 0.35 | 0.63 | 0.26 |
| S1HS_Mentally | 0.35 | 0.65 | 0.18 |
| S1HS_TiredEyes | 0.12 | 0.56 | 0.47 |
| S1VD1_OverDiff | 0.71 | 0.31 | 0.30 |
| S1VD1_Mentally | 0.71 | 0.31 | 0.24 |
| S1VD1_TiredEyes | 0.46 | 0.20 | 0.60 |
| S1FXS_OverDiff | 0.71 | 0.45 | 0.25 |
| S1FXS_Mentally | 0.70 | 0.45 | 0.26 |
| S1FXS_TiredEyes | 0.38 | 0.30 | 0.60 |
| S1RAN_OverDiff | 0.41 | 0.77 | 0.21 |
| S1RAN_Mentally | 0.40 | 0.76 | 0.20 |
| S1RAN_TiredEyes | 0.15 | 0.54 | 0.61 |
| S1TEX_OverDiff | 0.51 | 0.54 | 0.34 |
| S1TEX_Mentally | 0.37 | 0.56 | 0.38 |
| S1TEX_TiredEyes | 0.28 | 0.34 | 0.73 |
| S1BLG_OverDiff | 0.15 | 0.68 | 0.23 |
| S1BLG_Mentally | 0.27 | 0.69 | 0.27 |
| S1BLG_TiredEyes | 0.21 | 0.54 | 0.57 |
| S1VD2_OverDiff | 0.77 | 0.33 | 0.28 |
| S1VD2_Mentally | 0.76 | 0.35 | 0.27 |
| S1VD2_TiredEyes | 0.53 | 0.15 | 0.70 |
| S2HS_OverDiff | 0.43 | 0.72 | 0.23 |
| S2HS_Mentally | 0.47 | 0.72 | 0.18 |
| S2HS_TiredEyes | 0.17 | 0.54 | 0.59 |
| S2VD1_OverDiff | 0.81 | 0.33 | 0.32 |
| S2VD1_Mentally | 0.82 | 0.33 | 0.31 |
| S2VD1_TiredEyes | 0.57 | 0.15 | 0.71 |
| S2FXS_OverDiff | 0.68 | 0.46 | 0.32 |
| S2FXS_Mentally | 0.71 | 0.50 | 0.22 |
| S2FXS_TiredEyes | 0.43 | 0.28 | 0.67 |
| S2RAN_OverDiff | 0.39 | 0.73 | 0.31 |
| S2RAN_Mentally | 0.43 | 0.75 | 0.25 |
| S2RAN_TiredEyes | 0.14 | 0.49 | 0.70 |
| S2TEX_OverDiff | 0.50 | 0.52 | 0.46 |
| S2TEX_Mentally | 0.41 | 0.53 | 0.45 |
| S2TEX_TiredEyes | 0.29 | 0.25 | 0.82 |
| S2BLG_OverDiff | 0.28 | 0.62 | 0.31 |
| S2BLG_Mentally | 0.33 | 0.64 | 0.33 |
| S2BLG_TiredEyes | 0.20 | 0.45 | 0.67 |
| S2VD2_OverDiff | 0.78 | 0.29 | 0.36 |
| S2VD2_Mentally | 0.79 | 0.32 | 0.34 |
| S2VD2_TiredEyes | 0.50 | 0.11 | 0.75 |

Two items (S1TEX_OverDiff and S1TEX_Mentally) did not load cleanly on any factor (highest loading = .56) and were excluded from these factors. The reading task might not involve enough saccades to be grouped with tasks such as RAN and HS but it is not as easy as the free viewing task such as VD1 and FXS. Internal consistency estimates (Cronbach’s alpha) were .97 for Factor 1, .97 for Factor 2, and .96 for Factor 3.

### 4.3 Correlation between subjective reports

We conducted a correlation analysis between all subjective measurements including all session subjective reports, alcohol intake report, sleep duration report, caffeine intake report, and all reports after each task. This works as a sanity check for the data quality of these subjective reports. We expected the between-session and after-session fatigue to be correlated with the post-task reports, but not the before-session fatigue rating and post-task reports.

See figure this and that. The correlation results between all items from subjective reports (see Figure 7) as well as correlation between the major factors and averaged ratings (see Figure 7) showed consistent messages that subjective measures in GazeBase are consistent with each other, most likely reflecting the subjective fatigue of various aspects of the participants. Additionally, caffeine use, alcohol use, and sleep duration reports were included in the correlation analyses. Sleep showed a significant negative correlation with before-, between-, and after-session reports, but not the caffeine or alcohol intake. The correlation coefficient and statistical significance data can be seen in the appendix table.

**Figure 7:**
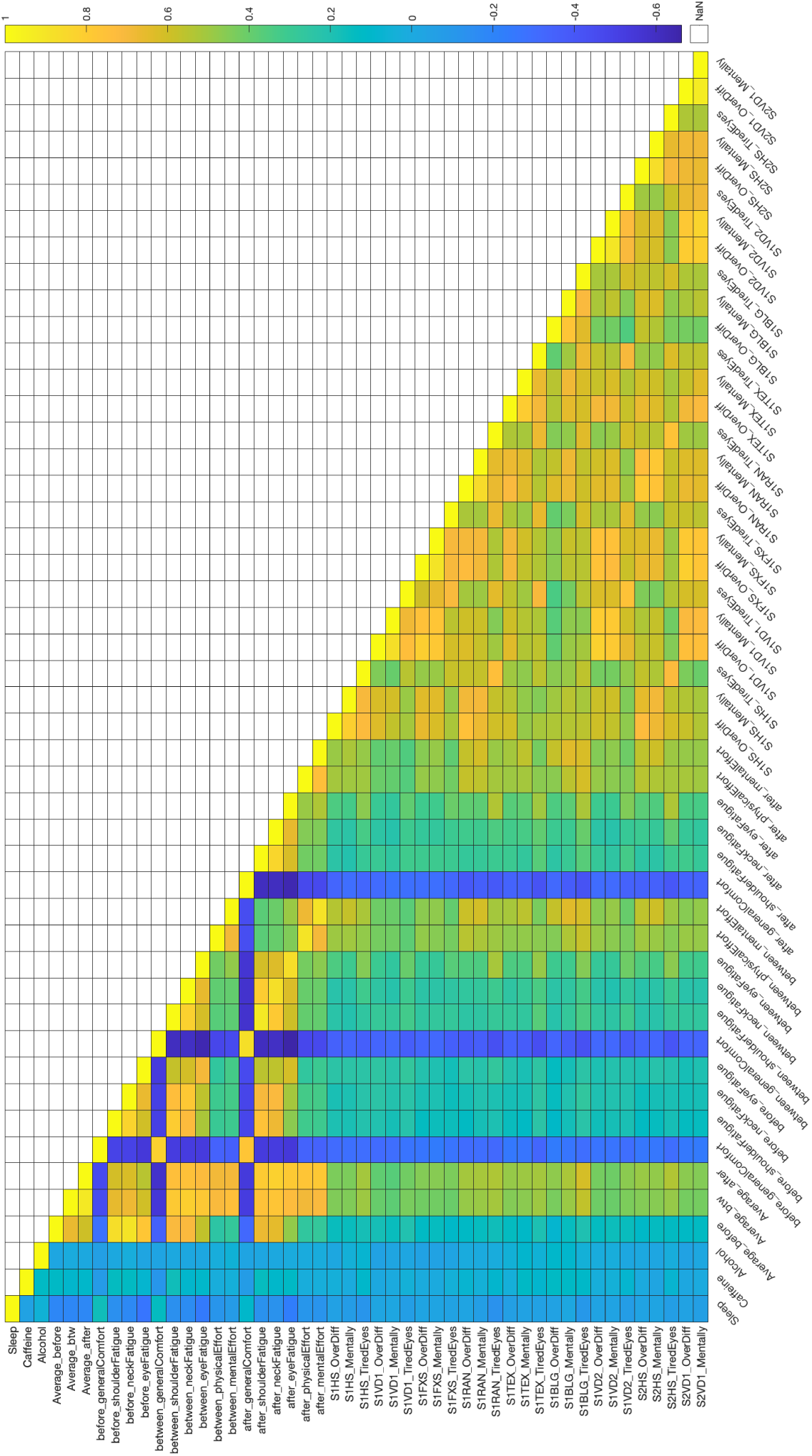
Correlation results between all subjective measurements.

**Figure 8:**
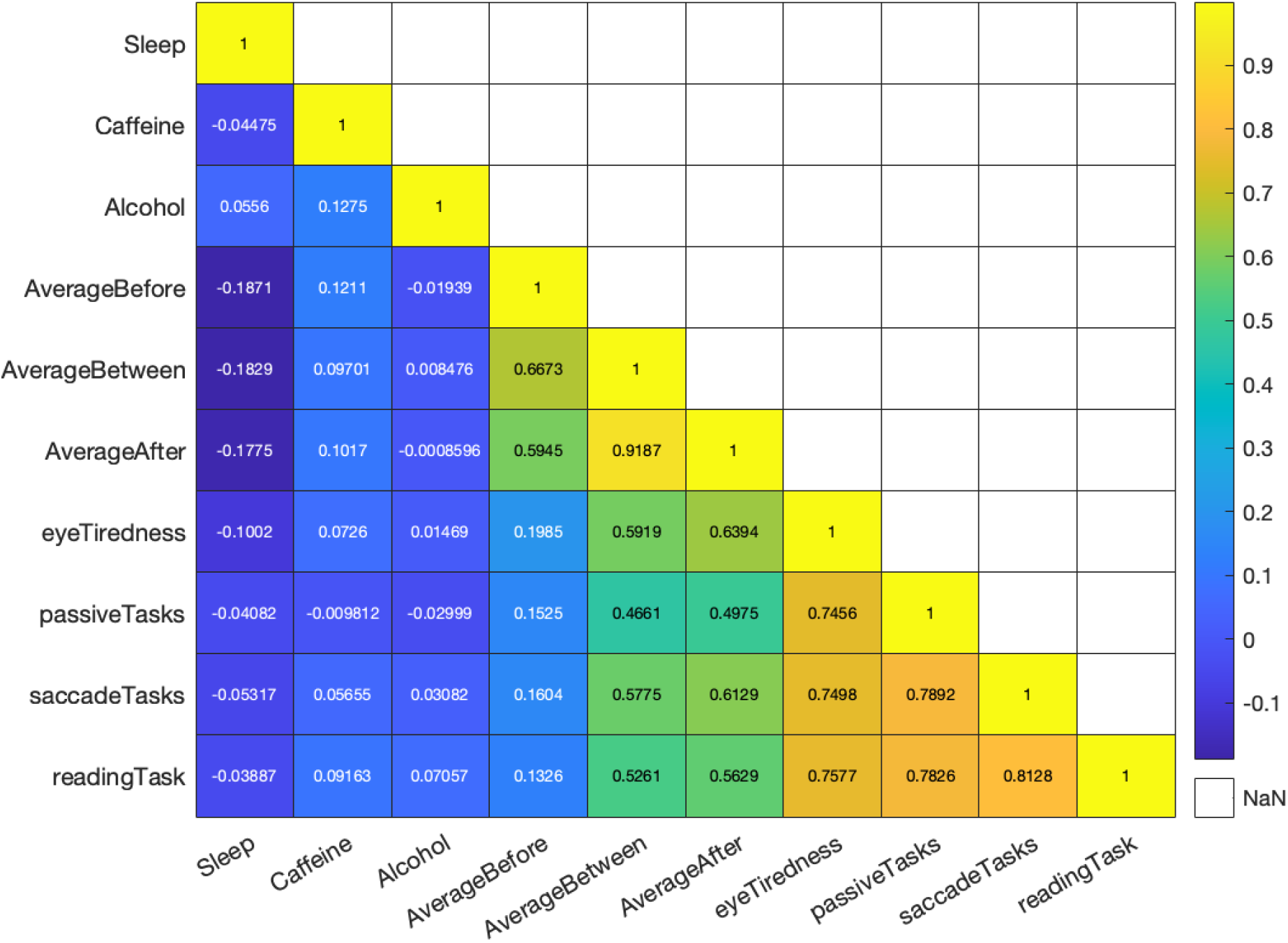
Correlation results between session subjective reports and EFA factors.

### 4.4 Validity demonstration with Balura game as an example

Gazebase is a dataset collected for the purpose of developing algorithms for identity identification from the perspective of data-driven approach. Unavoidably, the subjective report protocols were not evaluated or developed by experts in psychometrics. However, we hope to take advantage of the data as much as we can. It would be the best to evaluate the validity of every single task in this datasets, but the data suggested very little variance in many tasks such as fixation and video watching. Here, the task that reflected the largest amount of fatigue was evaluated.

Participants’ data were included if they produced subjective reports after completing the Balura game task, otherwise, their data are not included for this analysis. The resulting dataset contained 793 valid observations. We are treating two sessions completed approximately 15 minutes apart as separate observations. Participants who participated in this experiment multiple times in a longitudinal study manner (with at least one semester between rounds) were all included in this analysis.

We examined the relationship between subjective ratings and task duration by conducting Pearson’s correlation (see figure 9). Regarding three different subjective ratings (mental tiredness, subjective difficulty, and eye tiredness), they all showed a significant positive correlation with the duration of time spent on the BLG task and a transformed version of the duration (see statistics in the figure). We performed a linear fit on the median of each possible subjective rating and all of the fitting turned out to be significant, showing that the subjective reports, regardless of eye tiredness, mental tiredness, or difficulty rating, can predict the duration of BLG task completion, or vice versa. Further, this is a strong evidence suggesting the surveys participants completed indeed reflected their mental states. It simply requires a more difficult task to be more visible.

**Figure 9:**
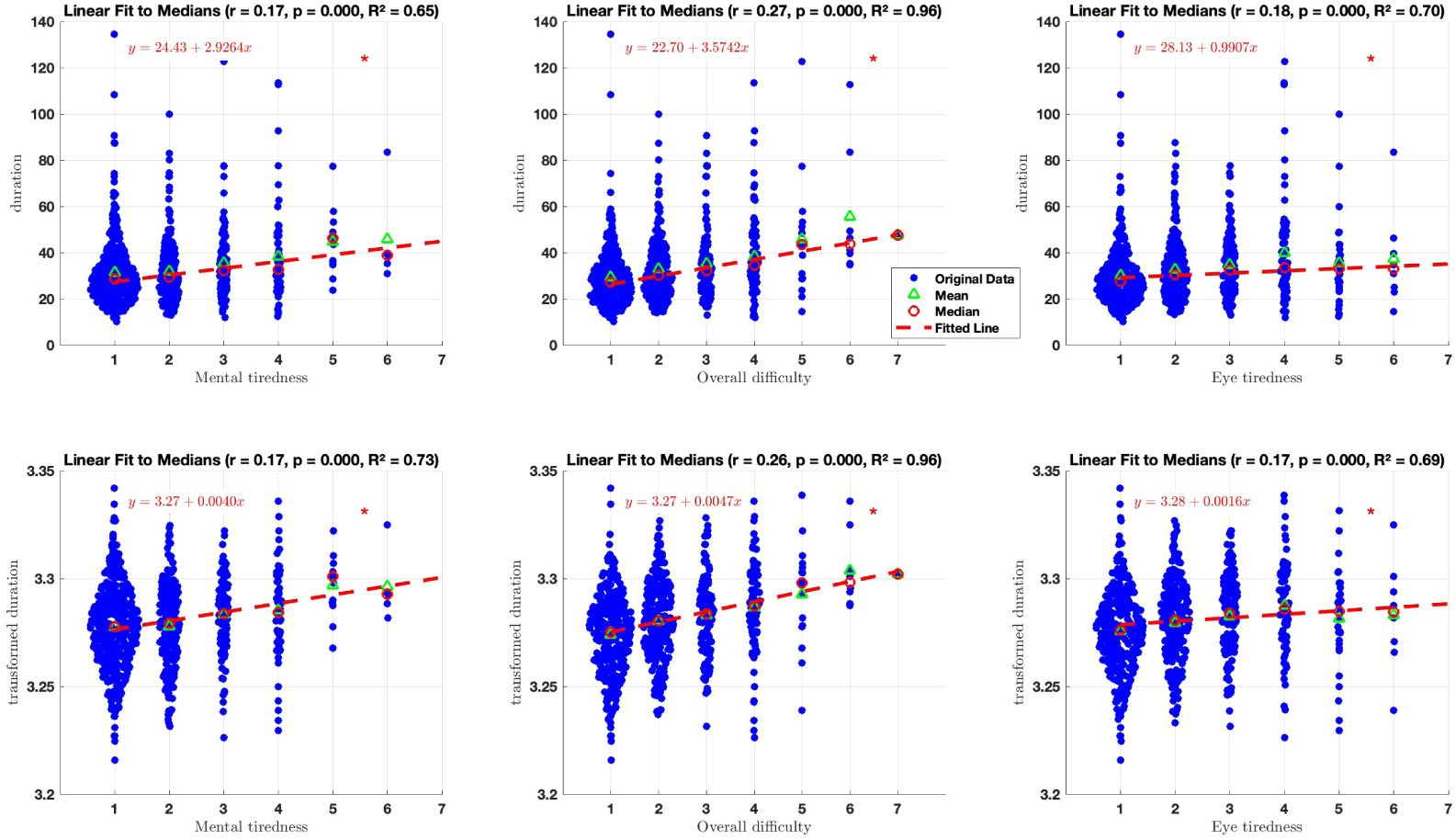
BLG task duration as a function of subjective ratings. The first row used the raw task duration in seconds and the second row used standardized duration data. The red dashed line is a linear fit of the median of the data. The red asterisk marks the significance of those fits.

We also fitted a few nonlinear curve fitting to understand the relationship between subjective reports and the duration on task completion. To model the relationship, four types of relationship were tested: linear, logarithmic (*y* = *a* + *b* ln(*x*)), exponential (*y* = *ae^bx^*), and power law (*y* = *ax^b^*). The goodness of fit for each model was evaluated using the coefficient of determination (*R*^2^), calculated as *R*^2^ = 1 − SS_res_*/*SS_tot_, where SS_res_ is the sum of squared residuals and SS_tot_ is the total sum of squares. To handle cases with near-constant dependent variables (variance < 10*^−^*^6^), *R*^2^ was set to 0 to avoid numerical instability. Except for the eye tiredness against the original duration, which had the best fit from the exponential model, the simple linear model produced the best fit for all figures.

### 4.5 Correlation between subjective reports and eye movement features

Linking subjective reports with objective eye movement measurements is one of the most straightforward ways to uncover state-specific or occasion-specific variances in eye movements. Here, we defined the subjective reports as a readout of the mental state. This is one of the motivations to obtain such datasets like GazeBase, and it is made available for other researchers. In this section, we correlated the subjective reports with features extracted from eye movements. Further, we examined the eye movement features that systematically change as a function of time both in the short term and in the long term and correlated these changes with subjective measurements.

#### 4.5.1 Eye movement features

Eye movement features were derived following the eye movement component identification [162, 163] and feature extraction protocol from Rigas et al. [164]. They organized features into two main categories: single-value features provide one value per recording, such as fixation rate or saccade rate, capturing overall temporal characteristics by dividing the total number of events by the recording duration; and multi-value features are derived from distributions of event-level measurements—such as fixation duration, saccade amplitude, or velocity—and summarized using descriptive statistics including the mean, median, standard deviation, inter-quartile range, skewness, and kurtosis. For our analysis, we retained all single-value features, as they represent overall temporal dynamics, and from the multi-value set, we selected only the median as a robust measure of central tendency, less affected by skewed distributions or outliers than the mean. This resulted in a focused set of features either the rate of certain eye movement component or median of the radius of a multi-value feature, consistent with the definitions and calculations established in the original study [164]. In total, we evaluated 58 features that can be found in the figure and tables below.

In this study, we deliberately prioritized features that are interpretable to human researchers, such as saccade rate and the median of drift acceleration, to facilitate better interpret the results and involve psychologists in the conversation. While other feature sets optimized for machine learning may offer greater statistical power through orthogonal or high-dimensional representations, such as features from [1], they often lack intuitive meaning, making it difficult to connect model performance with underlying physiological mechanisms. Our focus on interpretable features therefore supports reproducibility and cross-disciplinary communication, while leaving room for future work to integrate complementary, machine-learning–driven representations for enhanced predictive accuracy.

#### 4.5.2 Eye movement features and subjective reports

We conducted correlation analyses on three of the tasks: random saccades, horizontal saccades, and the reading task. The fixation task and video-watching tasks were easy for the participant, so they did not demonstrate enough variance for correlation analyses. Balura game is featured with pursuit eye movement and the existing feature extraction algorithm has not been optimized for pursuit yet. Considering that our subjective reports data do not conform to a normal distribution, we used Spearman correlation. The correlation results can be seen in Figures 10, 11, and 12. We acknowledge that these are not very strong effects and applying correction for multi-comparison will perish all of the significance.

**Figure 10:**
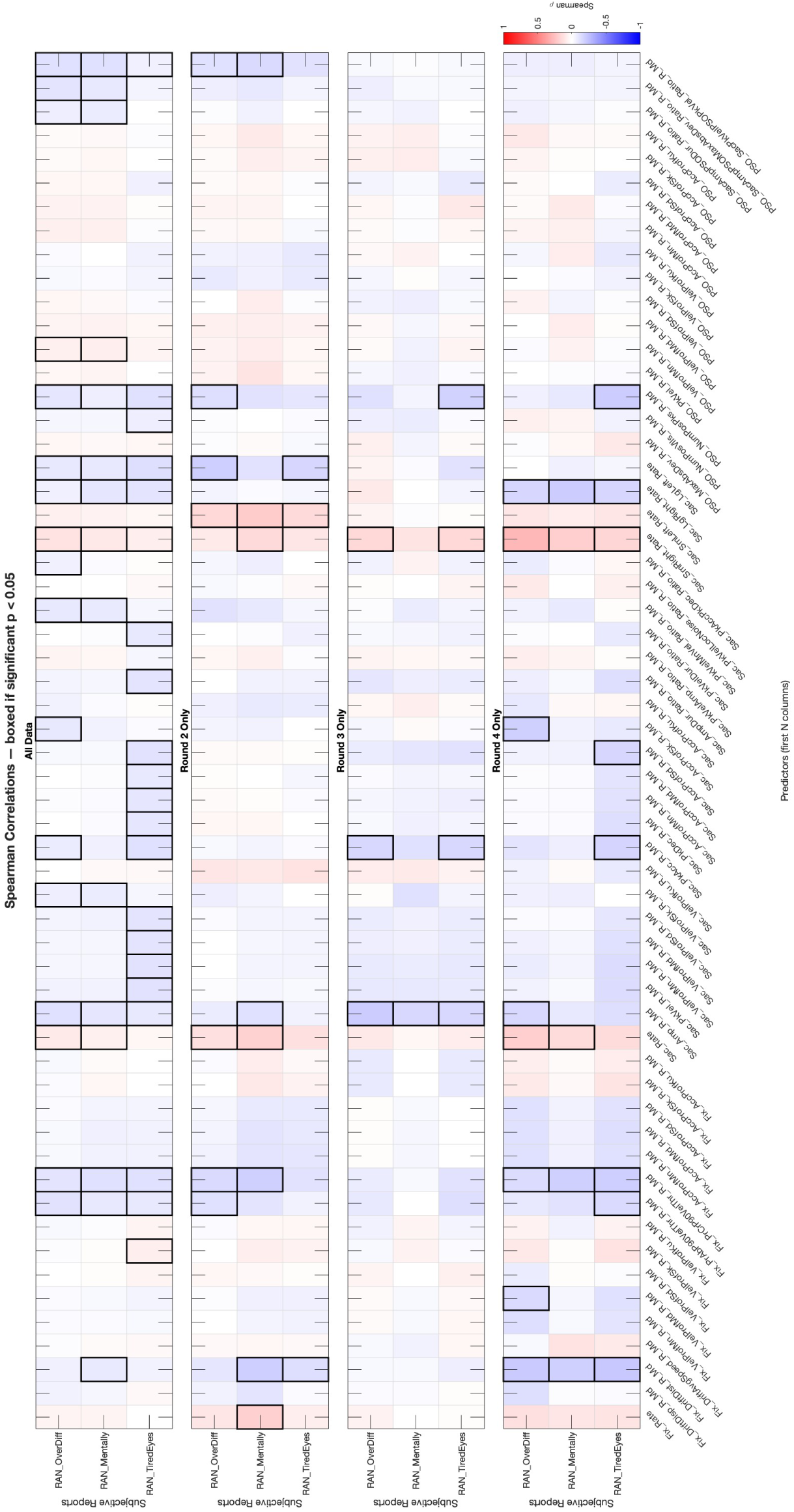
Correlation coefficients for subjective reports and eye movement features for random saccade task.

**Figure 11:**
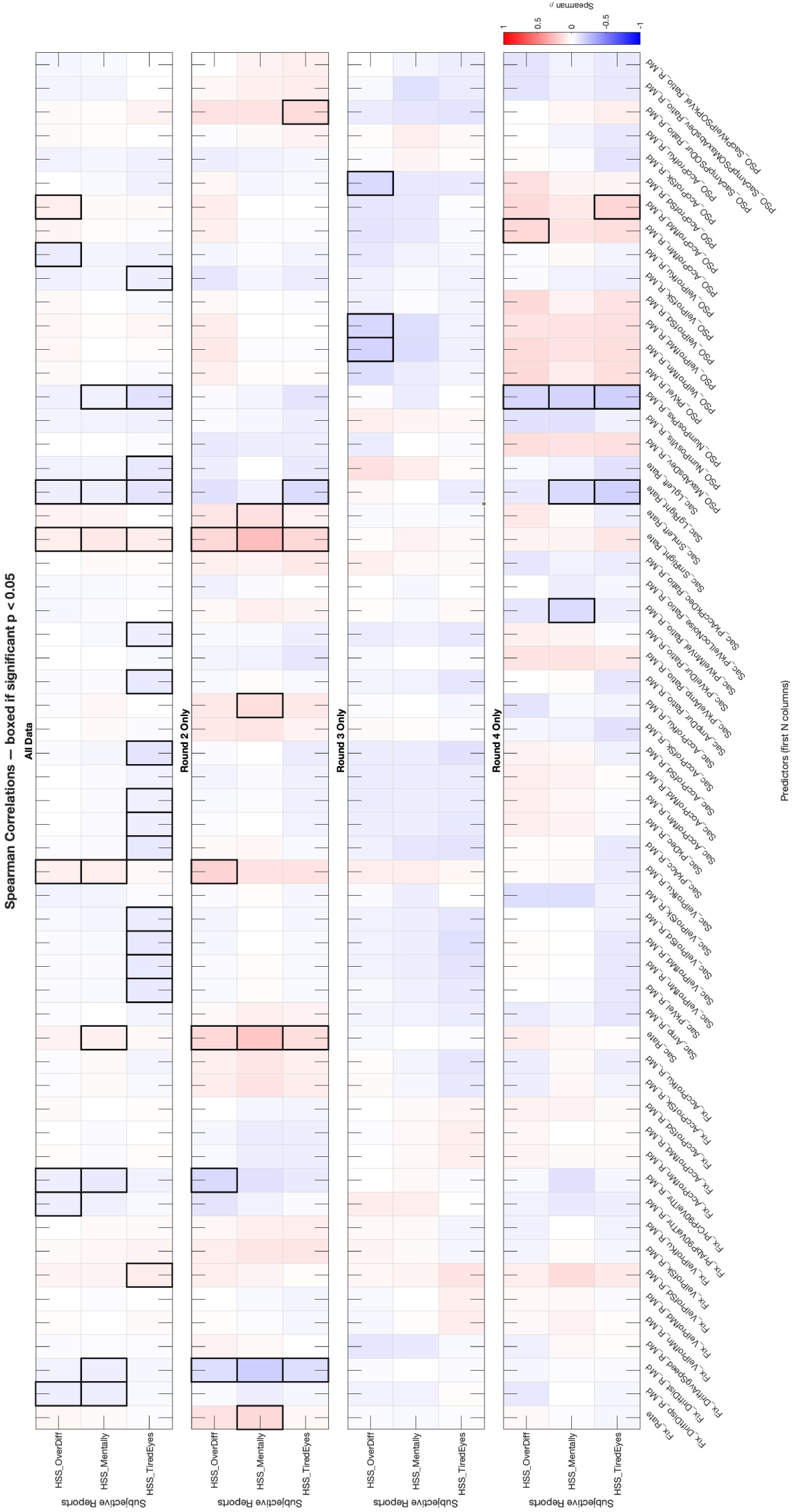
Correlation coefficients for subjective reports and eye movement features for horizontal saccade task.

**Figure 12:**
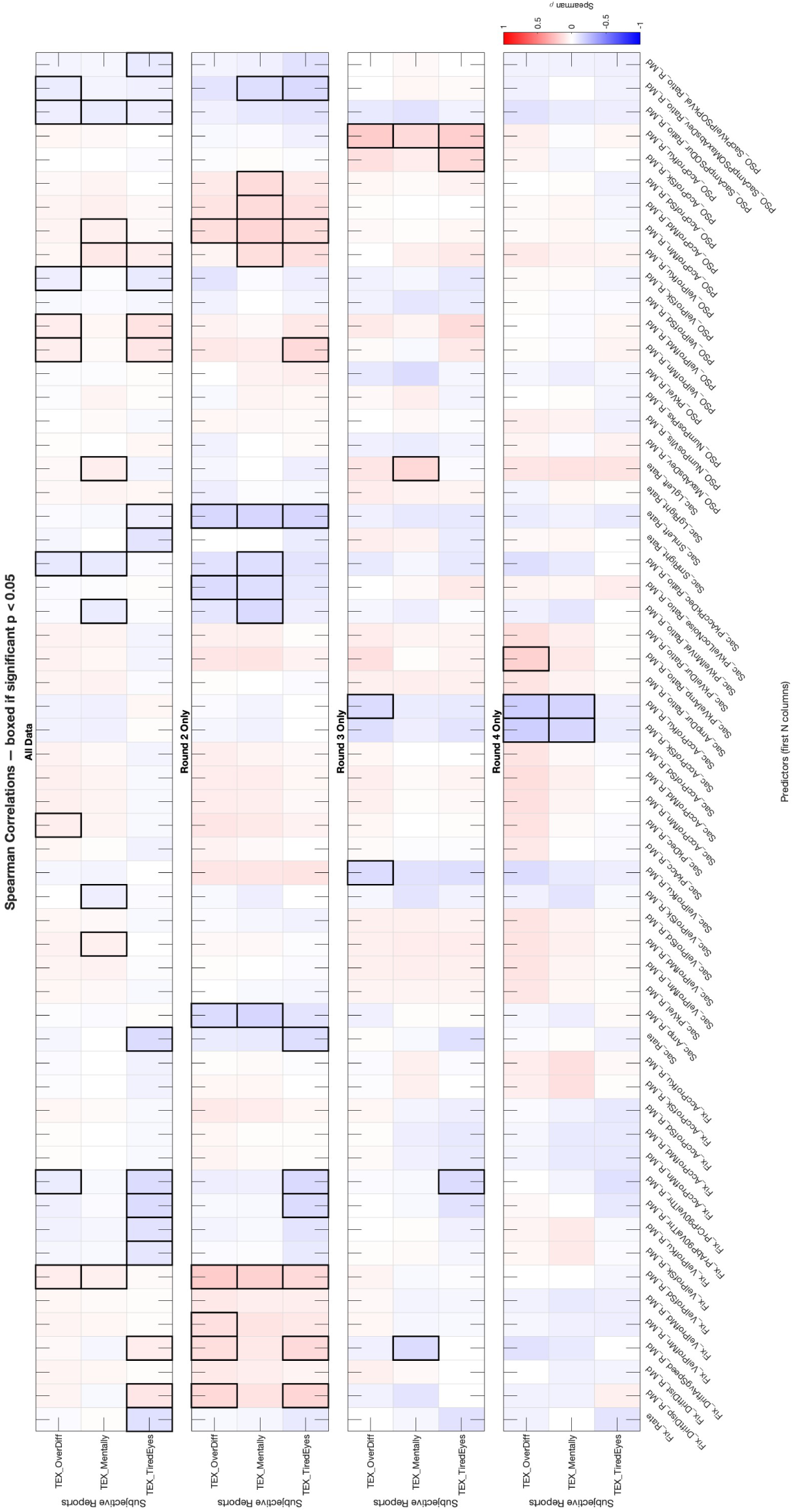
Correlation coefficients for subjective reports and eye movement features for reading task.

Across three tasks, we observed recurring associations between subjective reports and eye movement features related to saccade frequency and fixation stability. For instance, saccade rate correlated with subjective states in all three tasks, but with different emphases: in the random saccade task, higher saccade rates were positively correlated with eye tiredness reports; in the horizontal saccade task, they were positively correlated with mental tiredness; and in the reading task, the correlation reversed in direction with overall difficulty ratings. These effects were most consistently observed when combining all data or in Round 2, which had the most participants. In Rounds 3 and 4, the correlations were not consistently observed. This suggests that the two saccade tasks might share similar mechanisms. We speculate that if the reader found the reading task more difficult, they would take longer for each fixation and make fewer saccades as a result. For the saccade tasks, mental tiredness or eye tiredness were a result of higher saccade rates, which might be relevant to inaccurate saccade landings.

Fixation drift related features also showed correlations with subjective reports. The feature Fix_DriftDist_R_Md, captures the median radius of fixation drift distances, was correlated with eye tiredness and difficulty ratings across tasks. In the random saccade and horizontal saccade tasks, larger drift distances correlated negatively with reports of subjective tiredness, especially mental tiredness, indicating reduced fixation stability under fatigue. In the reading task, fixation drift distance did not show any significant correlations, but the drift average speed and drift discrepancy were positively correlated with eye tiredness and overall difficulty, but only in round 2. The differences in saccade and reading tasks contributed differently to the pattern of fatigue but showed instability during fixations when the participants experienced more fatigue; some are shown in drift distance and some are shown in speed or discrepancy. Similar to the correlations with saccade rate, these effects were most consistent when all data were combined or in Round 2 or occasionally in Round 4, whereas Rounds 3 and sometimes round 4 did not reproduce the same strength of correlation. These patterns point to fixation drift vigor as a potential describer of fatigue, though its expression appears sensitive to sample size and measurement round.

Together, the results for saccade rate and fixation drift vigor suggest that subjective fatigue states were reflected in eye movements. Although these patterns were not observed in every round of experiments, we judge this as a good starting point to further explore the eye movement features that can work as an index of fatigue. With machine learning method, these features can be combined and further information can be extracted and incorporated, making the signature of fatigue more pronounced. Further, the product developers might consider extracting the fatigue information for functions directly utilizing the fatigue meter, and developers can also work on removing the fatigue information in the data to achieve better privacy protection and privacy sharing controls.

Our results cannot identify the reasoning of the eye movement signature of fatigue. Future research can work on whether the fixation drift vigor is the cause of fatigue or whether it is a reflection of fatigue.

#### 4.5.3 Change of features over time

An intuitive way to index fatigue is to examine the trend of data as a function of time. We parsed the data from each session into 10 equal-sized chunks and used the same algorithm for feature extraction. Then we fitted a linear regression model and used the slope as an index of whether this feature changes as a function of time. See Figures 13, 14, and 15 for the features that demonstrate significant slope change as a function of time. Bonferroni correction was used for multiple comparisons.

**Figure 13:**
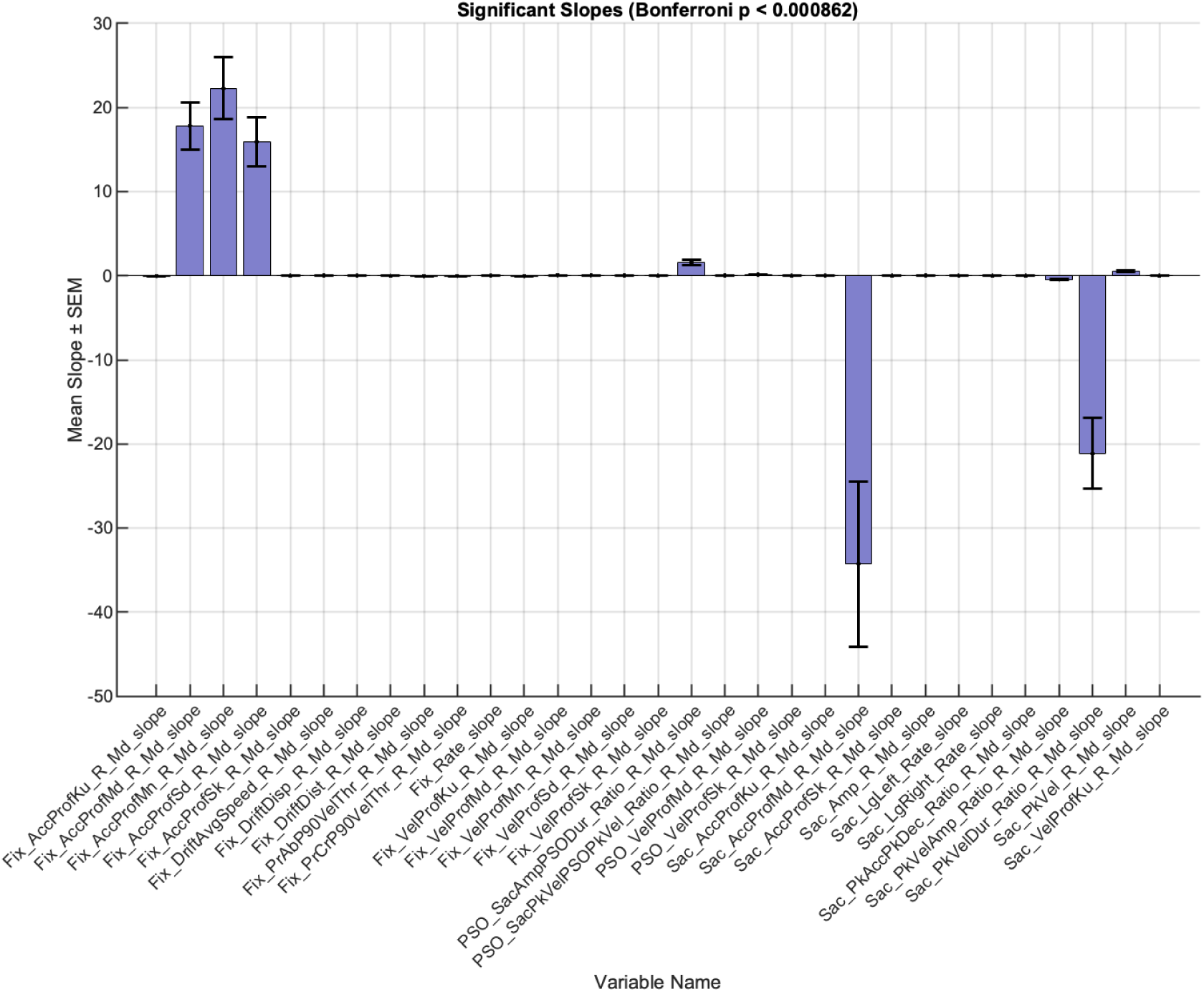
Significant slopes as a function of time for RAN task.

**Figure 14:**
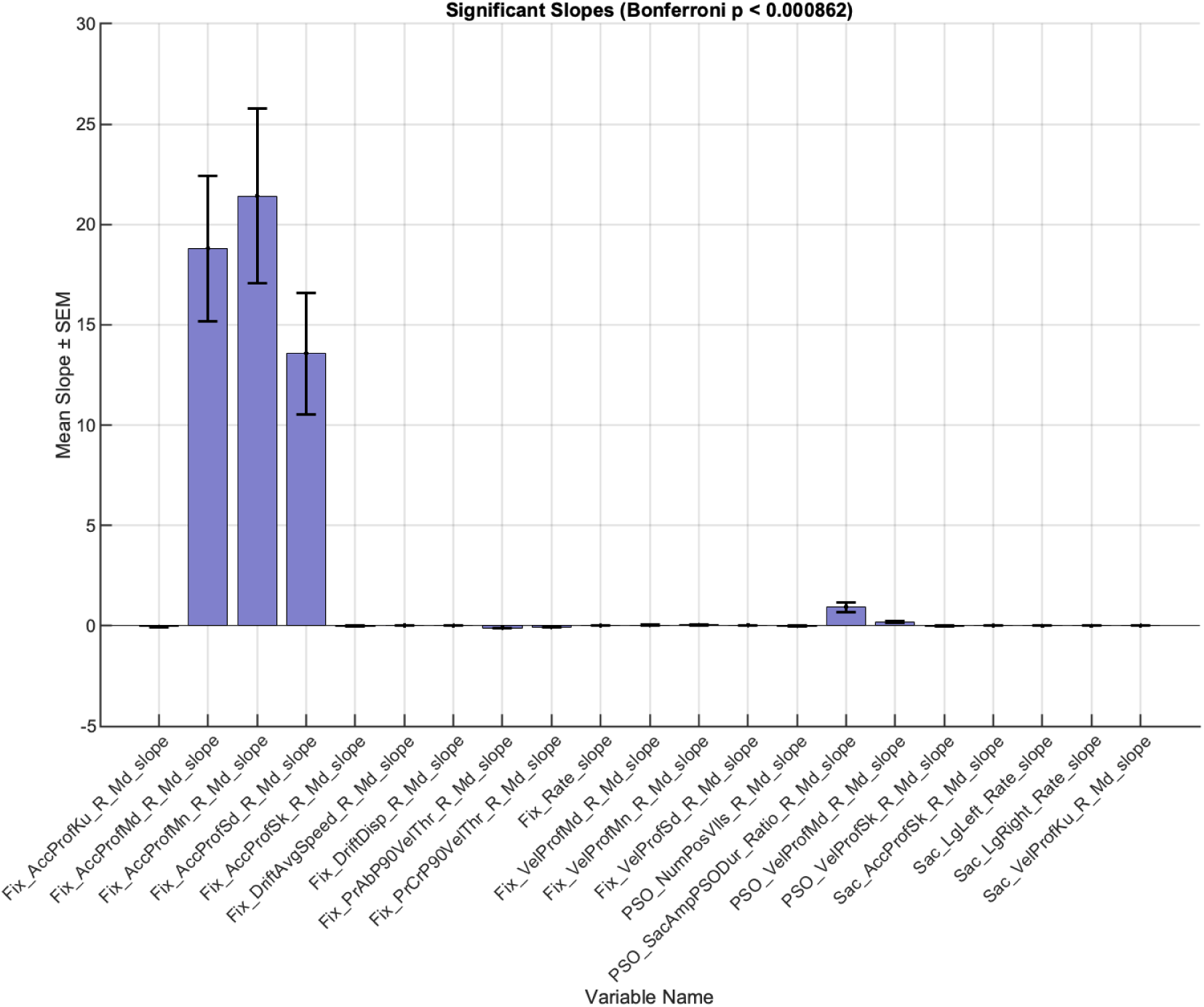
Significant slopes as a function of time for HSS task.

**Figure 15:**
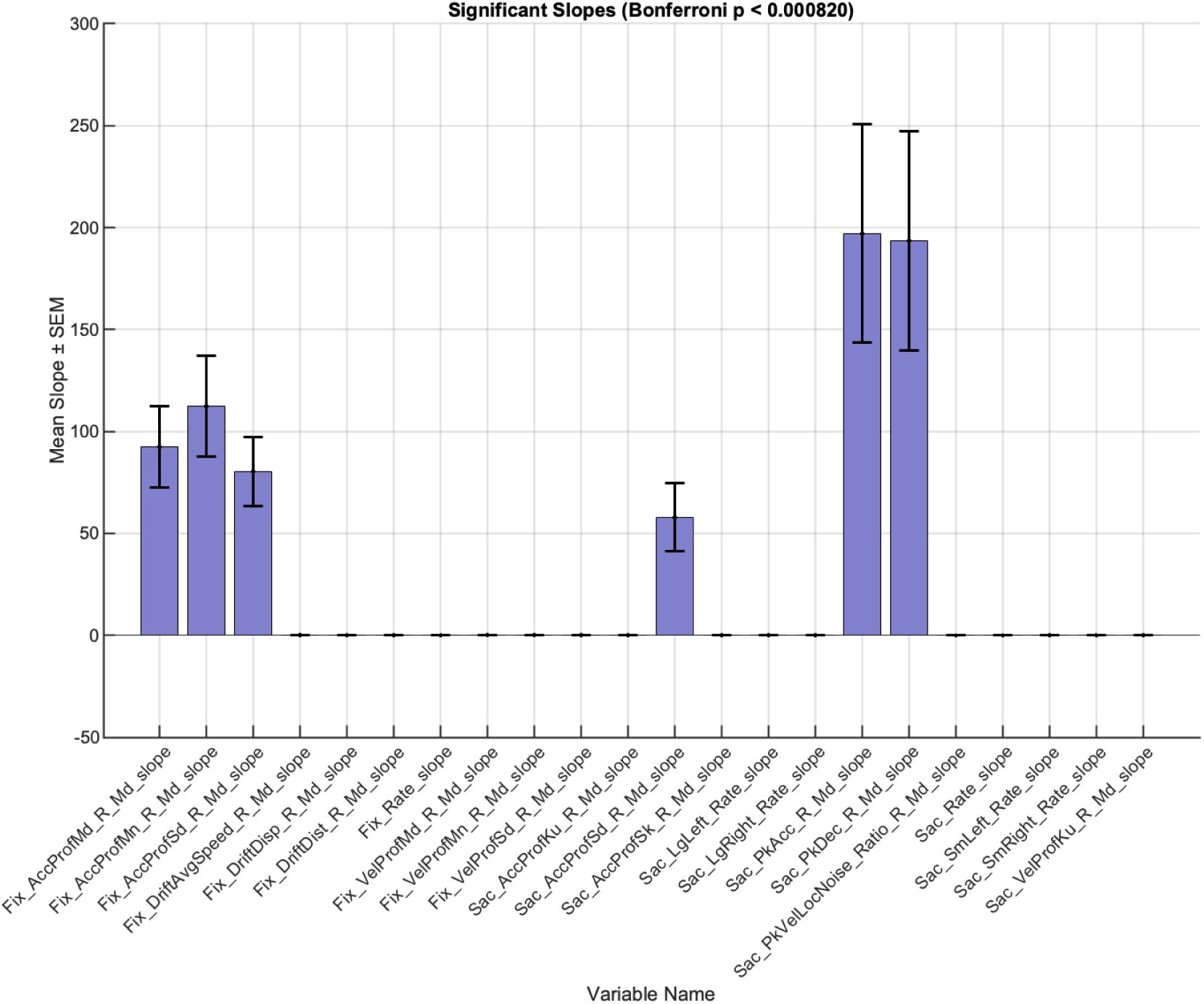
Significant slopes as a function of time for TEX task.

The analyses of the change of features as a function of time across three tasks drew our attention on acceleration profiles. Consistent between three tasks, features from the acceleration profile of fixations produced not only significant but also prominent positive mean slope as a function of time. This is a good candidate feature for indexing fatigue across tasks. For the reading task, peak accelerations and decelerations of saccades and the standard deviation of the acceleration profile of saccades increased over time. The median of the acceleration profile of saccades decreased over time for the RAN task. We also found a decrease of peak velocity-duration ratio of saccades over time for the RAN task. There are other features that reached significance in the model fit but the slope is too shallow to be interpretable.

We also correlated these slopes with subjective reports but did not observe any consistent results between rounds. It is likely that the tasks are too easy and too short for the healthy participants to feel fatigued enough, resulting in a measure that is not variable enough. It is also likely that the magnitude of these slopes is reflecting a very low level of change in fatigue that the participants cannot report it with the Likert measurement. Future research should test this with a longer and real time fatigue-indexed task.

## 5 Discussion

In this paper, we review the motivation for synthesizing oculomotor data and outline the theoretical and practical considerations that define how such data should look. When discussing why we need synthetic eye movement signals, we listed the limitations of existing real-world gaze datasets and the opportunities that synthetic data provide for scalability, privacy preservation, and reproducibility. In terms of actually synthesizing the signals, we identified core principles for realism and controllability, emphasizing task structure, inter-individual variability, and signal fidelity. Finally, the subjective reports data for GazeBase dataset is made available as training data for generating synthetic signals. We examine the relationship between self-reported subjective states and eye-movement features, showing a potential way to test synthetic signals. Together, this paper provides an integrated framework that connects conceptual motivation, design methodology, and empirical validation for advancing the development of synthetic oculomotor signals.

Engineers developing synthetic eye-movement data should ensure that the generated signals share the same fundamental components as real recordings, with comparable frequency distributions, durations, and other statistical characteristics. In addition, synthetic data should incorporate factors such as identity information, transient and sustained state variations, pipeline-specific differences, and realistic noise. These components can be systematically manipulated—for example, by removing identity cues while retaining state dynamics, or by aligning data from the same individual across different recording pipelines—to achieve specific design goals. Moreover, subjective reports and time-on-task measures can serve as proxies for internal state of fatigue, as they showed systematic variations in our analyses, and thus represent valuable dimensions for modeling and validation.

This dataset suggested a few aspects that future experimental studies can work on to better achieve other goals in data collection. 1) The order of the tasks can be shuffled with Latin square design [165] to minimize potential interference from different other tasks. 2) The subjective report data suggested a floor effect on the task difficulty. If fatigue is an important aspect of the study, we suggest to increase task difficulty such as adding an engaging task for passive viewing tasks, such as a preference task, counting task (gorilla study), or a description task. 3) Measurements before and after the task might help the researchers to get a better baseline of properties researchers are interested in. 4) Using a slide bar rather than a Likert scale can provide more information and make the transition between tasks and subjective reports more smoothly.

Our analyses of GazeBase dataset highlights several directions for improving future experimental designs to better support diverse goals in eye movement research or product development. Task order can be counterbalanced using a Latin square design [165] to minimize potential carryover or interference effects between tasks. The subjective report data also revealed a potential floor effect in perceived task difficulty, suggesting the need for more demanding or engaging activities—such as embedding preference judgments, object-counting tasks (as in the “gorilla” paradigm[166]), or open-ended description tasks within passive viewing conditions—to better capture fatigue and sustained attention. Baseline measurements collected both before and after task performance would provide a more stable reference for interpreting state-related changes in eye-movement features. Finally, implementing a continuous slider scale instead of a discrete Likert scale could yield more granularity in the subjective data.

The seemingly weak correlation or small effect size from psychologists’ perspective can turn into something quite useful from an engineering perspective. That the information being not accessible within the methods of traditional hypothesis testing or within a magnitude of 10 variables, does not mean such information can not be accessible at all. Here, a demonstration is the eye movement signature of individual differences. There are limited number of studies from psychologists’ perspective and the effects are solid but small. In contrast, the engineering approach already demonstrated successful identity recovery within a snippet of 5 seconds of sample reaching equal error rate of 3.66% accuracy [1].

A key distinction between synthetic face generation and the synthesis of eye-movement signals lies in how easily the realism of the output can be evaluated. Synthetic faces are immediately interpretable to humans—virtually everyone is a “face expert”—making perceptual validation straightforward. In contrast, synthetic eye-movement data, like other complex behavioral or physiological signals, cannot be meaningfully assessed through visual inspection alone and often require specialized analyses and domain expertise. This limitation underscores the need for interdisciplinary collaboration, where computer scientists work closely with psychologists, neuroscientists, and vision researchers to develop and validate generation evalulation procedures. Moreover, product developers should draw on the extensive, publicly available datasets collected in psychology and psychophysics, which capture rare or nuanced conditions often overlooked in large-scale datasets. Such insights go beyond the generation of synthetic eye movement data but other synthetic data.

## 6 Conclusions: Recommendations for Synthetic Data Generation

Real-world eye movement datasets remain expensive, logistically challenging, and constrained by privacy and diversity limitations. Synthetic eye movement data provide a practical and ethical alternative, but they should complement existing approaches. Datasets that lack full temporal resolution, such as fixation-only data, remain valuable for many research questions; synthetic data with richer temporal structure simply open new possibilities.

Future efforts in synthetic gaze generation should prioritize capturing the fine-grained temporal dynamics of natural eye movements—including fixations, saccades, blinks, pursuits, and microsaccades, while recognizing that simplified datasets tailored to specific needs continue to serve important roles. Synthetic data should also model both stable person-specific patterns and state-dependent changes such as fatigue or cognitive load, pipeline related differences, expanding their relevance for neuroscience, biometrics, and human–computer interaction. Crucially, behavioral metrics and subjective measures like task difficulty or fatigue should play a central role in guiding the design and validation of synthetic datasets, ensuring that they capture not only motor dynamics but also meaningful psychological states. Finally, the development of community benchmarks, open-source generation tools, and reproducible validation protocols will be essential for standardization, enabling fair comparisons between methods and ensuring ethical and responsible use.

By integrating these elements, the research community can build synthetic eye movement datasets that complement existing resources, incorporate behavioral dimensions, and overcome the cost, privacy, and scalability challenges of real-world data collection.

## 7 Acknowledgment

The data collection of GazeBase is funded by National Science Foundation grant CNS-1714623 to Dr. Komogortsev. The employment of C.S. Qian is supported by the school of Science and Engineering, Texas State University. We are grateful for the comments on statistical methods from our colleague, Dr. Lee Friedman, and we express our sincere condolences on his passing in August 2025.

