## Appendix for "Why do we need high-fidelity synthetic eye movement data and how should they look like?"

Nov 2025

### 1 Methods of GazeBase

In this appendix, we provide a detailed description of the experimental procedures and stimuli used to collect the GazeBase dataset [1]. The data is publicly available here and the code and additional data from this work is available here . Please refer to the original paper if you would like to know more details about the initiatives of the dataset. This dataset was intended to capture high-fidelity eye movement signals with emphases on longitudinal differences and between-subject differences across a battery of tasks. Participants were recruited from graduate and undergraduate students from Texas State University and they were incentivized to come for multiple rounds of data collection, with a minimum of 3-month gap. Each participant completed two recording sessions per laboratory visit, during which eye movements were captured across a fixed battery of seven tasks designed to elicit a broad range of oculomotor behaviors, from controlled saccades to naturalistic viewing and reading. Subjective self-report measures were collected at multiple points within each visit, and task-specific ratings were included in later rounds of data collection. The following subsections outline the design, purpose, and stimulus parameters of each task, along with notes on data collection protocols that are relevant for interpreting or reusing the dataset.

The data released beyond the previous release [1] is the subjective reports as well as more detailed demographics. Participants completed subjective self-report surveys at multiple points during each laboratory visit. Subjective reports were collected before the first session, between the two sessions, and after the final session. Participants used a Likert scale from 1 to 7 to rate general comfort, shoulder fatigue, neck fatigue, and eye fatigue at every reporting point. Additionally, they rated perceived physical and mental effort either between or after sessions. The initial questionnaire also included items about recent caffeine intake, sleep quality, alcohol consumption, and the presence of headaches. These survey instruments are publicly available with the present manuscript. Another package of the subjective reports started in the second round of data collection. Participants provided brief task-specific self-reports immediately following each experimental task. Using a 7-point Likert scale (1 = not at all; 7 = extremely), participants rated task difficulty, mental tiredness, and eye tiredness.

The experimenter instructed participants with a battery of tasks and performed 9-point calibration and validation before each individual task. The order of the tasks was kept the same. Participants were seated 55cm away from the monitor and they were instructed to stablize their chin and forehead on the chinrest.

### Task 1: Horizontal Saccade Task (HSS)

The HSS task elicited visually guided horizontal saccades of fixed amplitude by periodically shifting a peripheral target at 1-second intervals. Participants fixated the center of a bull's-eye target (red $1^\circ$ of visual angle enclosing a small black $0.5^\circ$ dot) identical to the calibration stimulus. The target appeared on a dark background (RGB value 0, 0 ,0) and was initially positioned at primary gaze location. It then alternated between positions at $\pm 15^\circ$ of visual angle, producing nominal $30^\circ$ horizontal saccades. Each position was held for 1s, for a total of 100 transitions per recording. Timing was constant across all participants. The experimenter verbally announced the remaining time in 20-second increments to help maintain engagement. Stimulus parameters and presentation order were identical across participants, sessions, and rounds.

### Task 2: Video Viewing Task 1 (VD1)

VD1 elicited naturalistic eye movements during free viewing of a video clip from a movie trailer. Participants watched the first 60 seconds of the trailer for *The Hobbit: The Desolation of Smaug* without audio. The same video segment was used across all participants and rounds. Due to variability across equipment configurations, the displayed duration during each second session was 57 s rather than 60 s.

### Task 3: Fixation Task (FXS)

The FXS task targeted fixational eye movements by presenting a static central bull's-eye fixation (same with the target in the HSS task) target for 15 seconds. Participants were asked to maintain fixation and, if possible, minimize blinking for the duration of the recording. The stimulus was identical for all participants, sessions, and rounds.

### Task 4: Random Saccade Task (RAN)

The RAN task elicited oblique, visually guided saccades of variable amplitude using a target that jumped to new positions every second. Participants followed the same bull's-eye fixation target used in earlier tasks. The target appeared on a dark background and moved to randomized screen locations spanning $\pm 15^\circ$ horizontally and $\pm 9^\circ$ vertically, with a minimum displacement of $2^\circ$ between consecutive locations. Each position remained visible for 1 s. Because target trajectories were randomized for each trial, stimulus sequences differed across participants, sessions, and rounds. The distribution of target locations ensured uniform spatial coverage. A sample screen-capture video from the RAN task is included in the data repository.

### Task 5: Reading Task (TEX)

The TEX task captured oculomotor behavior during reading. Participants silently read a 60-second segment of text drawn from *The Hunting of the Snark* by Lewis Carroll in font Caslon. The recording ended automatically after 60 seconds, regardless of reading progress. No specific instructions were provided for cases in which the participant finished early, although rereading the passage was suggested as one option. As a result, gaze patterns near the end of the recording may deviate from the typical line-by-line reading structure. Each session used a unique passage (18 total), with the same pair of passages assigned to all participants within a given round. Full-resolution images of all stimuli and documentation describing conversion from screen pixels to dva are available in the data repository.

### Task 6: Balura Game (BLG)

The BLG task measured eye movements during interaction with a gaze-controlled video game. On a black background, red and blue balls moved at a constant speed of $1^\circ$ per second. Initially, there are 10 red balls and 20 blue balls with a random spatial distribution across the whole screen. Participants attempted to eliminate all red balls as quickly as possible by fixating on them; blue balls could not be eliminated. A highlighted border appeared around a ball when a fixation was detected. The game ended once all red balls were removed. Further details are provided in `https://digital.library.txstate.edu/handle/10877/4158`, and a sample screen-capture video is included in the repository.

In some cases, sustained fixations on red balls did not trigger elimination. In such instances, the experimenter instructed participants to briefly look away and re-fixate on the ball. This behavior may appear as outward-and-return saccades in the recordings. Because ball positions and trajectories were randomized, stimuli differed across participants, sessions, and rounds.

### Task 7: Video Viewing Task 2 (VD2)

VD2 consisted of the subsequent 60 seconds of the same movie trailer used in VD1, also presented without audio. The same video file was used across participants and rounds. As with VD1, stimulus duration was shortened to 57 s in the second session of each visit due to technical constraints.
